# Phenotypic Noise and the Cost of Complexity

**DOI:** 10.1101/2020.02.26.963843

**Authors:** Charles Rocabert, Guillaume Beslon, Carole Knibbe, Samuel Bernard

## Abstract

Experimental studies demonstrate the existence of phenotypic diversity despite constant genotype and environment. Theoretical models based on a single phenotypic character predict that during an adaptation event, phenotypic noise should be positively selected far from the fitness optimum because it increases the fitness of the genotype, and then be selected against when the population reaches the optimum. It is suggested that because of this fitness gain, phenotypic noise should promote adaptive evolution. However, it is unclear how the selective advantage of phenotypic noise is linked to the rate of evolution, and whether any advantage would hold for more realistic, multi-dimensional phenotypes. Indeed, complex organisms suffer a cost of complexity, where beneficial mutations become rarer as the number of phenotypic characters increases. By using a quantitative genetics approach, we first show that for a one-dimensional phenotype, phenotypic noise promotes adaptive evolution on plateaus of positive fitness, independently from the direct selective advantage on fitness. Second, we show that for multi-dimensional phenotypes, phenotypic noise evolves to a low-dimensional configuration, with elevated noise in the direction of the fitness optimum. Such a dimensionality reduction of the phenotypic noise promotes adaptive evolution and numerical simulations show that it reduces the cost of complexity.

---

Isogenic populations of organisms often exhibit a diversity of phenotypes, even in constant environments. The importance of **phenotypic noise** (Yvert et al., 2013) has been highlighted with the development of experimental technologies measuring single individual variability (Elowitz et al., 2002; Raser and O’Shea, 2005; Ohya et al., 2015). Recent experimental studies indicate that phenotypic noise affects organismal fitness (Bódi et al., 2017; Duveau et al., 2018), is controlled genetically (Ordas et al., 2008; Hill and Mulder, 2010; Shen et al., 2012; Boukhibar and Barkoulas, 2016; Reyes et al., 2018) and is evolvable (Ito et al., 2009; Pelabon et al., 2010; Viñuelas et al., 2012; Shen et al., 2012; Metzger et al., 2015).

Phenotypic noise encompasses concepts such as *developmental noise* (Gavrilets and Hastings, 1994), *phenotypic heterogeneity* (Bódi et al., 2017), *cellular noise* (Hortsch and Kremling, 2018), *biological noise* (Eling et al., 2019), *intra-genotypic variability* (Bruijning et al., 2019). In quantitative genetics, phenotypic noise is historically known as **environmental variance** (or **micro-environmental variability**) (Falconer and Robertson, 1956). Environmental variance has long been assumed to be deleterious, because it decreases the **genotypic fitness** close to the fitness optimum and flattens the fitness landscape. On the other hand, Lande (1980) suggested that phenotypic noise should increase the genotypic fitness far from the fitness optimum. This phenomenon is due to the curvature of the fitness landscape, which is commonly assumed Gaussian. Close to the optimum, the fitness landscape is concave (its second derivative is negative, see Methods), and phenotypic noise decreases the genotypic fitness. Far from the optimum (beyond the inflection points, Pal 1998), the fitness landscape is convex (its second derivative is positive), and phenotypic noise increases the genotypic fitness (Pal, 1998; Kawecki, 2000; Paenke et al., 2007; Zhang et al., 2009; Bruijning et al., 2019). Hence, if the phenotypic noise level is evolvable, theoretical studies have shown that phenotypic noise should be positively selected in adaptive evolution because of its temporary positive effect on fitness (Pál and Miklós, 1999; Bruijning et al., 2019).

Since phenotypic noise increases the genotypic fitness of an organism far from the fitness optimum, it seems logical that it should facilitate adaptive evolution by increasing the fitness of beneficial mutations (Zhang et al., 2009; Bódi et al., 2017; Draghi, 2019; Schmutzer and Wagner, 2020). However, this claim is in contradiction with classical theory that states that phenotypic noise always reduces the **selection gradient** (Lande, 1975; Gavrilets and Hastings, 1994; Wang and Zhang, 2011; Mineta et al., 2015), as confirmed experimentally (Mulder et al., 2016; Keren et al., 2016). For example, essential genes and dosage-sensitive genes are under strong stabilizing selection and usually exhibit weak expression stochasticity (Fraser et al., 2004; Newman et al., 2006; Lehner, 2008). As suggested by Paenke et al. (2007), this confusion has two origins: **(i)** Historically, a Gaussian shape has been chosen to represent the fitness landscape because it “*has the merit of leading to algebraic simplicity* “(Robertson, 1956). **(ii)** There is a confusion between the impact of phenotypic noise on the genotypic fitness and its impact on the **gradient of relative genotypic fitness** (Paenke et al., 2007). Indeed, for Gaussian fitness functions, phenotypic noise can increase the genotypic fitness, but always decreases the gradient of relative genotypic fitness (Lande, 1976; Paenke et al., 2007). But apart for the algebraic simplicity, there is nothing generic in using a Gaussian fitness function. Finally, even if phenotypic noise increases the genotypic fitness for a one-dimensional phenotype, it is not clear how this generalizes to the level of an entire organism, which possesses “*phenotypically integrated complex units*” (Forsman, 2015). Fisher (1930) has suggested that organisms may pay a cost to the “complexity” of their phenotype, whereby the fraction of beneficial mutations becomes increasingly small when the number of phenotypic characters under selection increases. The cost of complexity hypothesis seems robust (Orr, 2000; Martin and Lenormand, 2006), and little affected by organismal modularity (Welch and Waxman, 2003) (but see Wagner et al. 2008). This raises questions about the effect of phenotypic noise when the number of phenotypic characters under selection increases: Will the advantage of phenotypic noise in a single dimension context will suffer from a cost of complexity?

Here, we look at the impact of an evolvable phenotypic noise on genotypic fitness and on the gradient of relative genotypic fitness in the context of **(i)** single-dimensional phenotypes, and **(ii)** multi-dimensional phenotypes. We first show how phenotypic noise impacts the adaptive evolution of a single phenotypic character towards a new fitness optimum. Using a quantitative genetics approach and a generalized fitness function, we show that, depending on the shape of the fitness function, phenotypic noise can increase the genotypic fitness of an organism, or promotes adaptive evolution, or both. We reveal the profound effect that the seemingly harmless addition of a minimal fitness to the fitness function can have on the evolutionary dynamics and on the evolutionary impact of phenotypic noise. We then extend the study to multiple phenotypic characters. We show that in an isotropic fitness landscape, an evolvable phenotypic noise is expected to align and correlate with the direction of the fitness optimum. Finally, using numerical simulations, we show that phenotypic noise can promote adaptive evolution of complex organisms and largely reduces the cost of complexity on the genotype when the phenotypic complexity is not too high.

## Methods

### Single phenotypic character under selection

We consider a large population of individuals under selection at discrete generations. We define an individual genotype by the pair {*µ, σ*}, where *µ* ∈ ℝ is the mean phenotype of the genotype {*µ, σ*} (or breeding value) and *σ* ∈ ℝ^+^ is the amplitude of the phenotypic noise. Both *µ* and *σ* are inheritable and evolvable. The phenotype *z* ∈ ℝ of a single organism with genotype {*µ, σ*} is assumed to be drawn from a normal distribution with mean *µ* and variance *σ*^2^: *p*(*z*|*µ, σ*^2^) ∼ 𝒩 (*µ, σ*^2^), but any distribution satisfying the Lindeberg’s condition can be used (Martin, 2014). The absolute fitness depends only on the phenotype *z*. The fitness function *w*(*z*) is assumed positive, three times differentiable, and has one non-degenerate optimum at *z* = 0 (Martin, 2014). We extended the fitness function from Tenaillon (2014) by adding a minimal fitness (Zhang et al., 2009; Draghi, 2019),

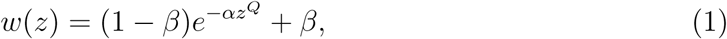

where *α >* 0 controls the sharpness of the fitness peak, *β* ∈ [0, 1] is the rescaled minimal fitness, and *Q* is a positive even integer controlling the curvature of the fitness function (Figure S1). When *β* = 0 and *Q* = 2, *w*(*z*) is Gaussian-shaped. Note that a non-null *β* is often used in numerical simulations (Zhang et al., 2009; Draghi, 2019).

The absolute fitness of the genotype {*µ, σ*} (or genotypic fitness) is (Lande, 1979):

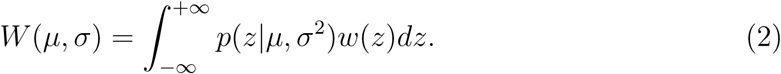

Analytical approaches to compute the effect of phenotypic noise on the genotypic fitness are detailed in Appendix S1. Numerical approaches are detailed in Script S1.

As discussed by Paenke et al. (2007), if all the genetic variation is additive or if reproduction is asexual, the response to selection equals the selection differential, *i*.*e*. the difference of the base population mean and the mean after selection (Lande, 1979). This does not hold with dominance or epistasis, but it is reasonable to assume that in most cases the response to selection will be positively correlated with the selection differential. Moreover, Paenke et al. (2007) have shown that if the effect of phenotypic noise on the genotypic fitness *W* (*µ, σ*) is monotonic, which holds when genetic variability is small, it is sufficient to study the impact of phenotypic noise on the gradient of relative genotypic fitness:

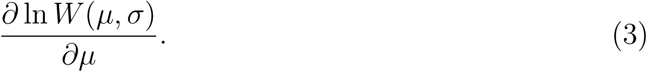

If 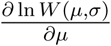 increases when *σ* increases, phenotypic noise magnifies the relative difference in fitness among genotypes and promotes adaptive evolution. Analytical approaches to compute the effect of noise on the gradient of relative genotypic fitness are detailed in Appendix S1. Numerical approaches are detailed in Script S1.

### Multiple phenotypic characters under selection

Fisher (1930) used the term “phenotypic complexity” to refer to the dimensionality *n* of the phenotypic space. Lande (1980) generalized quantitative equations for *n* phenotypic characters under selection. Here, we also included an evolvable multi-dimensional model of phenotypic noise.

In *n* dimensions, we define an individual genotype by the pair {***µ*, Σ**}, where ***µ*** = (*µ*_1_, *…, µ*_*n*_)^*T*^ ∈ ℝ^*n*^ is the mean phenotype (or breeding value) of the organism (*T* is the matrix transpose), and the *n* × *n* matrix **Σ** is the covariance matrix of the phenotypic noise. We assume that both ***µ*** and **Σ** are inheritable and evolvable. The continuous phenotype ***z*** = (*z*_1_, *…, z*_*n*_)^*T*^ ∈ ℝ^*n*^ follows a multivariate normal distribution centered at ***µ*** with covariance **Σ**: *p*(***z***|***µ*, Σ**) ∼ 𝒩_*n*_(***µ*, Σ**). We assume that the absolute fitness of an individual is determined by the Euclidean norm of its phenotype ‖ ***z*** ‖. Equation 1 generalizes to

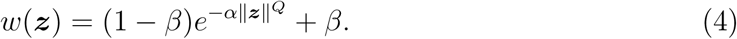

The fitness function in Equation 4 is **isotropic**, *i*.*e*. the fitness only depends on the Euclidean distance between the phenotype and the optimum. The absolute fitness of a genotype {***µ*, Σ**} (or genotypic fitness) is given by the multiple integral (Lande, 1980),

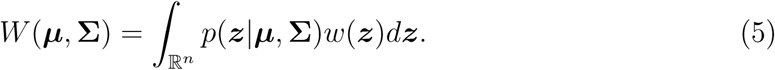

The *n* × *n* covariance matrix **Σ** is assumed real, symmetric and positive-definite. As such, it admits the eigenvalue decomposition

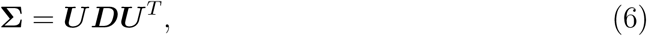

with a diagonal matrix ***D*** containing the *n* positive eigenvalues 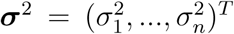 of **Σ**. The matrix ***U*** is a real orthonormal matrix that can be factorized as a product of rotations (***U*** can be chosen to avoid any reflections, see Anderson et al. 1987). **Σ** defines an hyper-ellipse in ℝ^*n*^ where the orientations of the semi-axes are given by the column vectors of ***U***, and the lengths of the semi-axes are given by the square roots of the eigenvalues (Fig. 1). This geometrical interpretation suggests a natural parametrization for the mutations; we express phenotypic noise mutations by **mutations in the lengths and in the orientations of the semi-axes of the hyper-ellipse**. Therefore, **Σ** is specified by a vector of *n* lengths ***σ*** = (*σ*_1_, *…, σ*_*n*_)^*T*^ and a vector of *n*(*n* – 1)*/*2 rotation angles ***θ*** = (*θ*_1_, *…, θ*_*n*(*n-*1)*/*2_)^*T*^. The genotype of an organism can be rewritten {***µ, σ, θ***}. The matrix ***U*** is built by applying successive rotations,

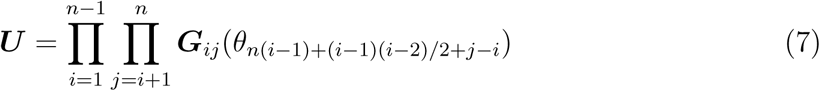

with ***G***_*ij*_(*θ*) the Givens matrix associated to the rotation between axes *i* and *j*, with an angle *θ*.

**Figure 1.**
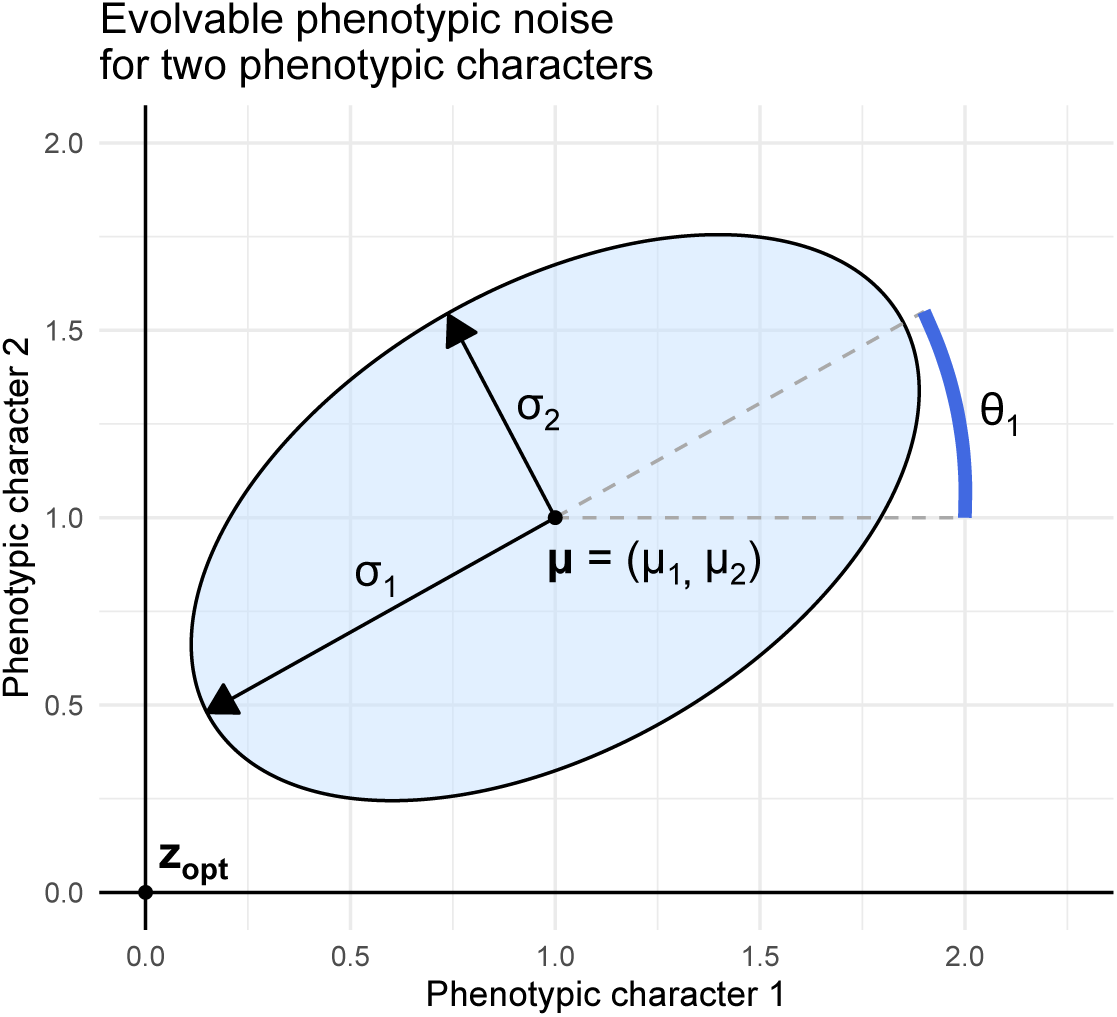
Phenotypic noise parametrization for two phenotypic characters under selection. For an organism with a mean phenotype ***µ*** = (*µ*_1_, *µ*_2_) (black dot in (1, 1)), phenotypic noise is defined by a vector of semi-axis lengths ***σ*** = (*σ*_1_, *σ*_2_) (black arrows) and a vector of rotations ***θ*** = (*θ*_1_) (for two phenotypic characters, a single rotation is needed, blue arc). The skyblue ellipse symbolizes the multivariate standard-deviation of the multivariate normal distribution of phenotypes for the genotype {***µ, σ, θ*}**. The fitness optimum ***z***_*opt*_ = (0, 0) is represented by the black dot at origin.

We assume that all coefficients of {***µ, σ, θ***} mutate independently. The vector of mutated mean phenotypes ***µ***^*′*^ is drawn from a multivariate normal distribution ***µ***^*′*^ ∼ 𝒩_*n*_(***µ, C***_***µ***_), the vector of mutated phenotypic noise amplitudes is drawn from a multivariate normal distribution ***σ***^*′*^ ∼ 𝒩_*n*_(***σ, C***_***σ***_), and the vector of mutated phenotypic noise orientations is drawn from a multivariate normal distribution ***θ***^*′*^ ∼ 𝒩_*n*(*n-*1)*/*2_(***θ, C***_***θ***_). ***C***_***µ***_, ***C***_***σ***_ and ***C***_***θ***_ are three constant covariance matrices of size *n* × *n* for ***C***_***µ***_ and ***C***_***σ***_, and of size *n*(*n* – 1)*/*2 × *n*(*n* – 1)*/*2 for ***C***_***θ***_. In addition to the mutational variability, we assign mutation probabilities per generation *m*_*µ*_, *m*_*σ*_ and *m*_*θ*_. The mutational distributions have finite mean and variance and satisfy the Lindeberg’s condition (Martin, 2014).

In summary, our model includes three evolvable vectors per individual: ***µ, σ*** and ***θ***. The mean phenotype of each organism is represented by a vector ***µ***. The phenotypic noise of each organism is modeled by a multivariate normal law 𝒩_*n*_(***µ*, Σ**), **Σ** being decomposed in its semi-axes of lengths ***σ***, and rotation angles ***θ***. The model also includes three constant mutational covariances ***C***_***µ***_, ***C***_***σ***_ and ***C***_***θ***_, three constant mutation rates *m*_*µ*_, *m*_*σ*_ and *m*_*θ*_ (per individual per generation), and an isotropic fitness function *w*(***z***) (Eq. 4). Table 1 summarizes all the variables.

**Table 1.** List of mathematical variables.

| Variable notation | Type | Description |
| --- | --- | --- |
| • Single phenotypic character under selection |  |  |
| $z$ | $\mathbb{R}$ | Phenotype quantitative value |
| $w(z)$ | $\mathbb{R} \rightarrow \mathbb{R}$ | Fitness function (Eq. 1) |
| $\alpha$ | $\mathbb{R}^+$ | Sharpness of the fitness peak |
| $\beta$ | $[0, 1]$ | Minimal fitness value |
| $Q$ | $2k, k \in \mathbb{N}^*$ | Curvature of the fitness function |
| $z_{opt}$ | $\mathbb{R}$ | Optimal phenotype |
| $\mu$ | $\mathbb{R}$ | Mean phenotype (or breeding value) |
| $\sigma$ | $\mathbb{R}$ | Phenotypic noise standard deviation |
| $W(\mu, \sigma)$ | $\mathbb{R} \times \mathbb{R} \rightarrow \mathbb{R}$ | Genotypic fitness given $\mu$ and $\sigma$ (Eq. 2) |
| $\partial W(\mu, \sigma)/\partial \mu$ | $\mathbb{R} \times \mathbb{R} \rightarrow \mathbb{R}$ | Genotypic fitness gradient given $\mu$ and $\sigma$ |
| $\partial \ln W(\mu, \sigma)/\partial \mu$ | $\mathbb{R} \times \mathbb{R} \rightarrow \mathbb{R}$ | Relative genotypic fitness gradient given $\mu$ and $\sigma$ |
| $\partial W(\mu, \sigma)/\partial \sigma$ | $\mathbb{R} \times \mathbb{R} \rightarrow \mathbb{R}$ | Genotypic fitness gradient around $\sigma$, given $\mu$ |
| $\partial^2 \ln W(\mu, \sigma)/\partial \mu \partial \sigma$ | $\mathbb{R} \times \mathbb{R} \rightarrow \mathbb{R}$ | Relative genotypic fitness gradient around $\sigma$, given $\mu$ |
| • Multiple phenotypic characters under selection |  |  |
| $n$ | $\mathbb{N}^*$ | Dimensionality of the phenotypic space (“complexity”) |
| $\mathbf{z}$ | $\mathbb{R}^n$ | Phenotype quantitative value |
| $w(\mathbf{z})$ | $\mathbb{R}^n \rightarrow \mathbb{R}$ | Fitness function (Eq. 4) |
| $d$ | $\mathbb{R}^n \times \mathbb{R}^n \rightarrow \mathbb{R}$ | Euclidean distance to the optimal phenotype |
| $\mathbf{z}_{opt}$ | $\mathbb{R}^n$ | Optimal phenotype |
| $\boldsymbol{\mu}$ | $\mathbb{R}^n$ | Mean phenotype (or breeding value) |
| $\boldsymbol{\Sigma}$ | $\mathbb{R}^{n \times n}$ | Covariance matrix of the phenotypic noise |
| $W(\boldsymbol{\mu}, \boldsymbol{\Sigma})$ | $\mathbb{R}^n \times \mathbb{R}^{n \times n} \rightarrow \mathbb{R}$ | Genotypic fitness given the genotype $\{\boldsymbol{\mu}, \boldsymbol{\Sigma}\}$ (Eq. 5) |
| $\mathbf{D}$ | $\mathbb{R}^{n \times n}$ | Diagonal matrix of the eigenvalues of $\boldsymbol{\Sigma}$ |
| $\mathbf{U}$ | $\mathbb{R}^{n \times n}$ | Real orthogonal matrix of the eigenvectors of $\boldsymbol{\Sigma}$ |
| $\sigma^2$ | $\mathbb{R}^n$ | Eigenvalues of $\boldsymbol{\Sigma}$ |
| $\boldsymbol{\theta}$ | $\mathbb{R}^{n(n-1)/2}$ | Pairwise plane rotations to generate $\mathbf{U}$ |
| $\mathbf{C}_\mu$ | $\mathbb{R}^{n \times n}$ | Mutational covariance matrix of $\boldsymbol{\mu}$ |
| $\mathbf{C}_\sigma$ | $\mathbb{R}^{n \times n}$ | Mutational covariance matrix of $\boldsymbol{\sigma}$ |
| $\mathbf{C}_\theta$ | $\mathbb{R}^{n(n-1)/2 \times n(n-1)/2}$ | Mutational covariance matrix of $\boldsymbol{\theta}$ |
| $\mathbf{G}_{ij}(\theta)$ | $\mathbb{R}^{n \times n}$ | Givens matrix (rotation between axes $i$ and $j$, with angle $\theta$) |
| $s_\mu^2$ | $\mathbb{R}^+$ | Mutational variance of $\boldsymbol{\mu}$ ($\mathbf{C}_\mu = s_\mu^2 \mathbf{I}_n$) |
| $s_\sigma^2$ | $\mathbb{R}^+$ | Mutational variance of $\boldsymbol{\Sigma}$ ($\mathbf{C}_\sigma = s_\sigma^2 \mathbf{I}_n$) |
| $s_\theta^2$ | $\mathbb{R}^+$ | Mutational variance of $\boldsymbol{\theta}$ ($\mathbf{C}_\theta = s_\theta^2 \mathbf{I}_{n(n-1)/2}$) |
| $\mathbf{I}_n$ | $\mathbb{R}^{n \times n}$ | Identity matrix of size $n \times n$ |
| $m_\mu$ | $[0, 1]$ | Mutation rate of $\boldsymbol{\mu}$ |
| $m_\sigma$ | $[0, 1]$ | Mutation rate of $\boldsymbol{\Sigma}$ |
| $m_\theta$ | $[0, 1]$ | Mutation rate of $\boldsymbol{\theta}$ |
| $L$ | $\mathbb{N}^*$ | Population size |

### Numerical simulations

We simulated the adaptive evolution of a population of individuals undergoing evolvable phenotypic noise under stabilizing selection. We considered a population of constant size *L*. At each discrete generation, individuals reproduce asexually with mutations. The number of descendants depends on the relative fitness and genetic drift. Each descendant inherits a mutated genotype {***µ***^*′*^, ***σ***^*′*^, ***θ***^*′*^} from its parents {***µ, σ, θ***}, depending on the constant mutational covariances ***C***_***µ***_, ***C***_***σ***_ and ***C***_***θ***_, and the constant mutation rates *m*_*µ*_, *m*_*σ*_ and *m*_*θ*_. We made the additional assumption that mutation sizes were isotropic for ***µ, σ*** and ***θ***. Mean traits *µ*_*i*_, *i* ∈ {1, *…, n*} independently mutate through a normal distribution 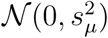. Each semi-axis size *σ*_*i*_, *i* ∈ {1, *…, n*} independently mutates through a normal distribution 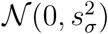. Rotation angles *θ*_*i*_, *i* ∈ {1, *…, n*(*n* – 1)*/*2} independently mutate through a normal distribution 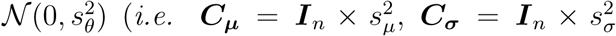 and 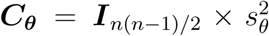). We used three different measures of evolution; assuming *σ*_1_ = max(***σ***):

i. The **phenotypic noise amplitude** *σ*_1_, *i*.*e*. the square root of the principal eigenvalue, which is the standard deviation of the phenotypic noise along the principal axis of **Σ**.
ii. The **phenotypic noise flattening**,

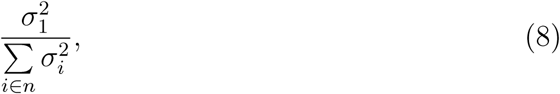

which is the fraction of the total variance along the principal axis. If the flattening ≈1, all the variance is contained in the principal axis of the phenotypic noise (*i.e*. there is a dimensionality reduction of phenotypic noise towards a single dimension).
iii. The **phenotypic noise alignment**,

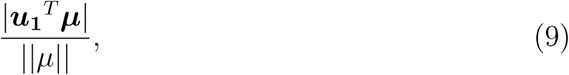

which is the correlation between the eigenvector associated to the largest eigenvalue, and the direction of the optimum. If the correlation ≈ 1, the principal axis of the phenotypic noise is aligned towards the optimum.

For each numerical simulation, both the population’s mean and the population’s variance of these measures were computed through time. This model extends Fisher’s geometric model (Fisher, 1930) by adding an evolvable phenotypic noise. The code of the simulation framework is available in Script S2. Table 1 summarizes all the variables.

## Results

### For a single phenotypic character, phenotypic noise promotes adaptive evolution on plateaus of positive fitness

Standard results (Pal, 1998) show that non-null phenotypic noise is beneficial to the genotypic fitness (Eq. 2) beyond the inflection points of the fitness function and deleterious inside. There is an optimal noise amplitude (achieved when 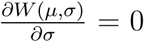) due to the smoothing effect at larger noise levels (Fig. 2, *solid black lines*). A classic result with Gaussian fitness functions states that phenotypic noise always decreases the gradient of relative genotypic fitness (Lande, 1975). However, phenotypic noise can promote adaptive evolution in other kinds of fitness functions (when *β* ≥ 0 and/or *Q* ≥ 2, Eq. 1). We show in Appendix S1 that phenotypic noise promotes adaptive evolution on plateaus of positive fitness (*i.e*. on log-fitness plateaus). When a population is located on such a plateau where the selection gradient is negligible, phenotypic noise increases this gradient by magnifying the relative genotypic fitness differences present at the borders of the plateau, hence favoring the fixation of beneficial mutations (see Appendix S1). In Equation 1, the formation of plateaus of positive fitness depends on the minimal fitness *β* and the fitness curvature *Q*, and never occurs in the Gaussian setting. Indeed, if the fitness function is based on Equation 1, the log-fitness plateaus only:

**Figure 2.**
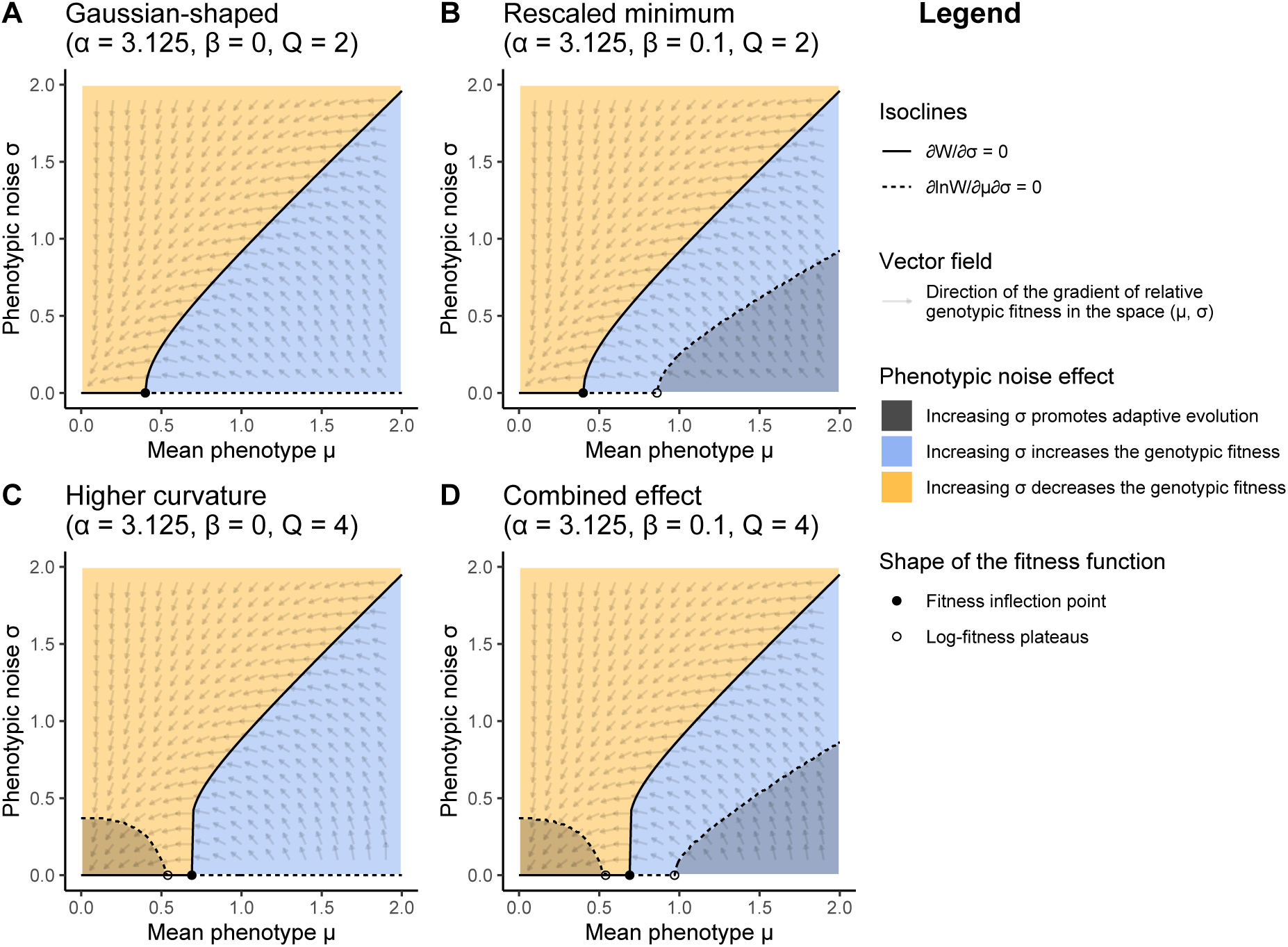
Impact of phenotypic noise on the genotypic fitness and the gradient of relative genotypic fitness, depending on the shape of the fitness function. **A**. Gaussian-shaped fitness landscape (*α* = 3.125, *β* = 0 and *Q* = 2). **B**. Rescaled minimum fitness landscape (*α* = 3.125, *β* = 0.1 and *Q* = 2). **C**. Higher curvature fitness landscape (*α* = 3.125, *β* = 0 and *Q* = 4). **D**. Combined effects fitness landscape (*α* = 3.125, *β* = 0.1 and *Q* = 4). The blue and orange areas represent the regions where increasing *σ* respectively increases or decreases the genotypic fitness *W* (*µ, σ*). The grey area represents the region where increasing *σ* increases the gradient of relative genotypic fitness 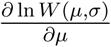, hence promoting adaptive evolution. The isocline 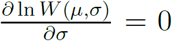 for which *σ* maximizes *W* (*µ, σ*) is represented by a solid black line. The isocline 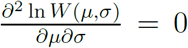 for which *σ* maximizes 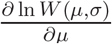 is represented by a dashed black line. Fitness function inflection point is represented by a black dot. The distance below or beyond which the fitness function plateaus is represented by black circles. The direction of the gradient of relative genotypic fitness is represented by a vector field in the space (*µ, σ*).

i. far from the optimum when *β >* 0,
ii. around the optimum when *Q >* 2.

The exact threshold values of the mean phenotype *µ* below or beyond which phenotypic noise promotes adaptive evolution on fitness plateaus are provided in Table 2.

**Table 2.**
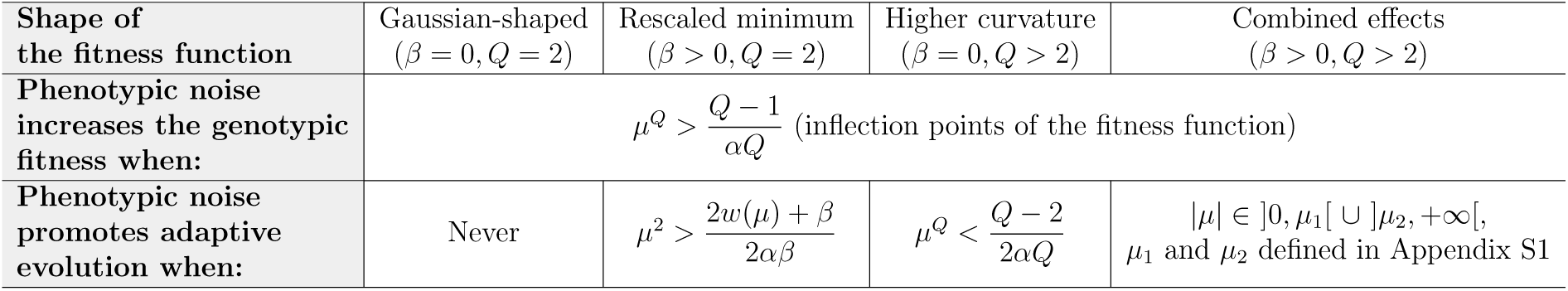
Threshold values of the mean phenotype *µ* beyond or below which phenotypic noise increases the genotypic fitness or promotes adaptive evolution, depending on the shape of the fitness landscape.

In summary, during periods of adaptation to new environments, phenotypic noise is expected to be positively selected far from the fitness optimum because it increases the genotypic fitness. It will also promote adaptive evolution if the log-fitness plateaus far from the optimum (*i.e*. if *β >* 0). This should be followed by a reduction in phenotypic noise when the population reaches the optimum, because phenotypic noise decreases the genotypic fitness, even if it promote adaptive evolution if the log-fitness plateaus around the optimum (*i.e*. if *Q >* 2). Thus, while phenotypic noise has the potential to indirectly speed up the fixation of beneficial mutations far and/or close to the fitness optimum depending on the shape of the fitness landscape, its actual role will depend on how direct selection impacts phenotypic noise based on its effect on the genotypic fitness. This raises the question of the indirect selection of phenotypic noise for its specific impact on the gradient of relative genotypic fitness (see Discussion).

To illustrate this result, we took as a reference the parameters used by Zhang et al. (2009) (*α* = 3.125, *β* = 0.1 and *Q* = 2). We then selected four configurations of fitness functions, by varying the minimal fitness *β* and the fitness curvature *Q* (*α* plays no role here).

1. Gaussian-shaped fitness function (*α* = 3.125, *β* = 0 and *Q* = 2) (Fig. 2A),
2. Rescaled minimum fitness function (*α* = 3.125, *β* = 0.1 and *Q* = 2) (Fig. 2B),
3. Increased curvature fitness function (*α* = 3.125, *β* = 0 and *Q* = 4) (Fig. 2C),
4. “Combined effects” fitness function (*α* = 3.125, *β* = 0.1 and *Q* = 4) (Fig. 2D).

For simplicity, we only explored positive values of the mean phenotype *µ*, as the result is symmetrical around *z*_*opt*_ = 0. In all configurations, the phenotypic noise increases the genotypic fitness *W* (*µ, σ*) in convex regions of the fitness function, beyond the inflection point (Fig. 2 *black dots*). The optimal *σ* value maximizing *W* (*µ, σ*) increases as the distance between the mean phenotype *µ* and the optimum increases (Fig. 2 *solid black lines*). As expected, for a Gaussian fitness function (Fig. 2A), the phenotypic noise always decreases the gradient of relative genotypic fitness, *i.e*. it slows down adaptive evolution (Lande, 1975). When the minimal fitness *β >* 0 (*i.e*. the log-fitness plateaus at its minimum ln(*β*), Fig. 2B *black circle*), phenotypic noise promotes adaptive evolution but only far from the optimum. This mechanism is independent from the curvature of the fitness landscape, and does not depend on the inflection point (Fig. 2B *black dot*). The optimal *σ* value maximizing the gradient of relative genotypic fitness increases as the distance between *µ* and the optimum increases (Fig. 2B *dashed black line*). When the fitness curvature parameter *Q* increases, a log-fitness plateau appears around the optimum (Fig. 2C *black circle*), and phenotypic noise promotes adaptive evolution. The *σ* value maximizing the gradient of relative genotypic fitness decreases quickly as the distance between *µ* and the optimum increases (Fig. 2C *dashed black line*). Therefore phenotypic noise can promote adaptive evolution both close and far from the optimum (Fig. 2D *dashed black line*).

The impact of phenotypic noise on the genotypic fitness and the gradient of relative genotypic fitness is summarized in Table 3. During an adaptation event, according to the vector field

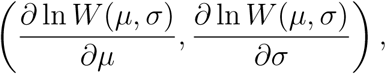

an evolvable *σ* is expected to increase far from the optimum, and then to decrease to zero as the population reaches the fitness optimum (Fig. 2 *vector field*).

**Table 3.**
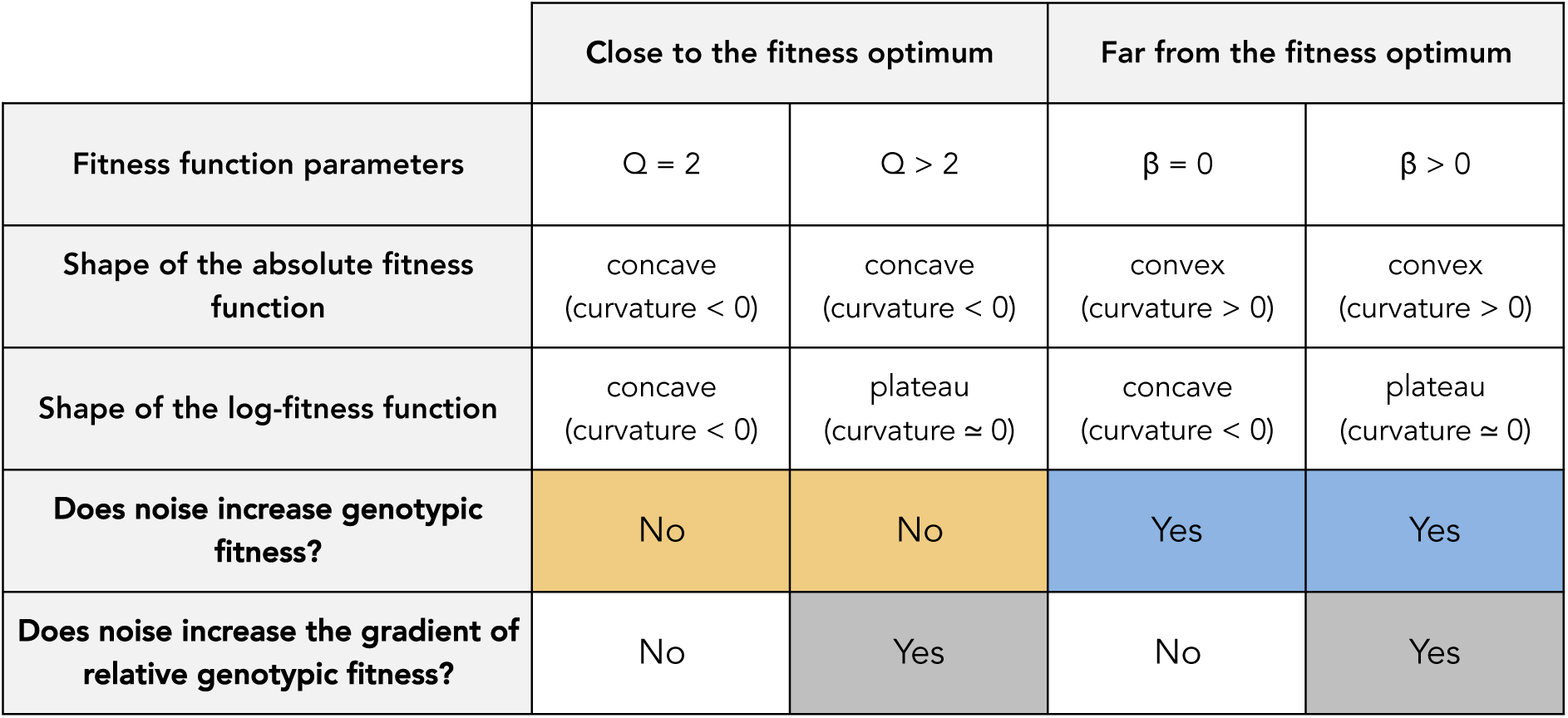
Impact of phenotypic noise on the genotypic fitness and the gradient of relative genotypic fitness, depending on the shape of the fitness function. Orange, blue and dark-grey background colors correspond to the color code used in figure 2.

|  | Close to the fitness optimum |  | Far from the fitness optimum |  |
| --- | --- | --- | --- | --- |
| Fitness function parameters | $Q = 2$ | $Q > 2$ | $\beta = 0$ | $\beta > 0$ |
| Shape of the absolute fitness function | concave<br>(curvature $< 0$) | concave<br>(curvature $< 0$) | convex<br>(curvature $> 0$) | convex<br>(curvature $> 0$) |
| Shape of the log-fitness function | concave<br>(curvature $< 0$) | plateau<br>(curvature $\approx 0$) | concave<br>(curvature $< 0$) | plateau<br>(curvature $\approx 0$) |
| Does noise increase genotypic fitness? | No | No | Yes | Yes |
| Does noise increase the gradient of relative genotypic fitness? | No | Yes | No | Yes |

For each fitness function, we also used numerical simulations to compute evolutionary trajectories (Methods, Script S2), in order to estimate the convergence time towards the fitness optimum. The mean phenotype mutation size was *s*_*µ*_ = 0.1, and its mutation rate was *m*_*µ*_ = 10^−3^ (∼1 mutation per generation per population). For a single phenotypic character, there is no possible rotation of the single phenotypic noise component, thus ***θ*** = ∅ (Methods). We ran simulations with a population size *L* = 1, 000 for the four fitness configurations when there is:

1. no phenotypic noise (*σ* = 0),
2. an evolvable phenotypic noise with a mutation size *s*_*σ*_ = 0.1 and a mutation rate *m*_*σ*_ = 10^−3^ (∼1 mutation per generation per population).

A population was assumed to reach the fitness optimum when the mean phenotype in the population was at a distance 0.1 from the optimum (one quarter of the fitness standard deviation). We ran 200 repetitions for each of the 8 parameter sets (1,600 simulations in all). All individuals were initialized with the genotype {*µ* = 2, *σ* = 0}. When phenotypic noise promotes adaptive evolution far from the optimum (*i.e*. when the log-fitness plateaus; *β >* 0), populations with an evolvable noise converge faster to the optimum (Fig. 3B-D). When phenotypic noise does not promote adaptive evolution far from the optimum (*i.e*. when *β* = 0, *e.g*. for a Gaussian-shaped fitness function), populations without phenotypic noise converge faster (Figs. 3A-C). In all the cases where phenotypic noise is evolvable, genotypes {*µ, σ*} tend to follow the gradient of relative genotypic fitness in the space (*µ, σ*) to maximize the genotypic fitness (Figure S2). Doing so, the population may go through regions of the space (*µ, σ*) where the phenotypic noise promotes adaptive evolution, indirectly reducing the convergence time depending on the shape of the fitness function.

**Figure 3.**
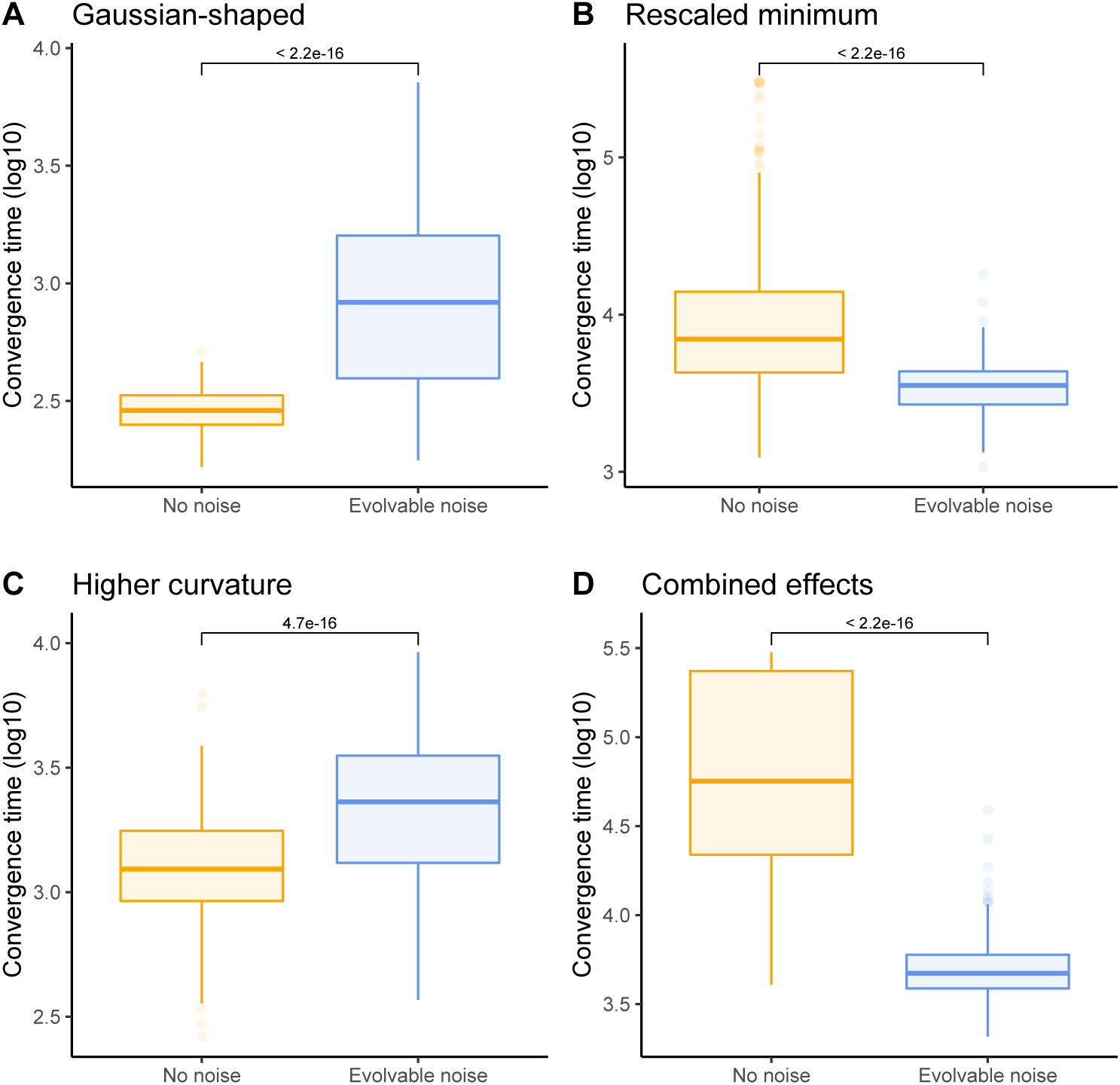
Population convergence time to the fitness optimum depending on the shape of the fitness function and the evolvability of phenotypic noise. **A**. Gaussian-shaped fitness landscape (*α* = 3.125, *β* = 0 and *Q* = 2). **B**. Rescaled minimum fitness landscape (*α* = 3.125, *β* = 0.1 and *Q* = 2). **C**. Higher curvature fitness landscape (*α* = 3.125, *β* = 0 and *Q* = 4). **D**. Combined effects fitness landscape (*α* = 3.125, *β* = 0.1 and *Q* = 4). The population with no phenotypic noise (*σ* = 0) is represented by an orange boxplot. The population with evolvable phenotypic noise is represented by a blue boxplot. Wilcoxon test p-values are provided.

### For multiple phenotypic characters, the best phenotypic noise configuration is aligned and fully correlated with the direction of the fitness optimum

When organisms have several phenotypic characters under selection, the fraction of beneficial mutations decreases quickly with the number *n* of characters, a property known as the “cost of complexity” (Fisher, 1930; Orr, 2005). This effect is maximized under the “universal pleiotropy assumption” (Martin, 2014), where mutations on the mean phenotype are isotropic, such that mutations have no preferential direction and can affect all characters similarly. When genetic correlations between phenotypic characters exist and are evolvable, the universal pleiotropy does not hold anymore (see *e.g*. Schluter 1996; Sato and Kaneko 2019), reducing the cost of complexity. Here, we keep the universal pleiotropy assumption as a worse-case scenario, in order to properly characterize the role of an evolvable phenotypic noise in complex phenotypes.

We show in Appendix S1 that when the population is far from the fitness optimum (beyond the inflection points of the fitness landscape), the best phenotypic noise configuration (*i.e*., the one that maximizes the genotypic fitness) is reached when phenotypic noise is fully correlated and aligned with the direction of the fitness optimum. In this configuration, the phenotypic noise is reduced to a single dimension, with an optimal value in the direction of the fitness optimum and no noise in all other directions. Any other form of phenotypic noise yields a lower genotypic fitness *W* (***µ*, Σ**) (Eq. 5). Thus, the best phenotypic noise configuration for *n* phenotypic characters far from the fitness optimum consists in a dimensionality reduction to fight the cost of complexity on phenotypic noise. A population evolving such a noise configuration will recover the benefit of a single character scenario, with phenotypic noise conferring a strong fitness advantage to organisms. However, the effect of phenotypic noise on the gradient of relative genotypic fitness will still depend on the shape of the fitness landscape, at least in the sub-space where phenotypic noise generates random phenotypes. Inside the inflection points of the fitness landscape, the best configuration is a null phenotypic noise.

An example of such an evolvable phenotypic noise is shown in Figure 4. In this numerical simulation (see Methods, Script S2), a population of individuals with two phenotypic characters evolve towards the fitness optimum. As shown on Figure 4A, phenotypic noise shape evolves through generations and eventually reach the optimal configuration, fully flattened and aligned with the fitness optimum located in (0, 0). As the mean phenotype in the population converges (Fig. 4B), the phenotypic noise amplitude increases temporarily (Fig. 4C). Phenotypic noise is also temporarily reduced to a single dimension, as shown by the flattening of the phenotypic noise (Fig. 4D). At the same time, phenotypic noise temporarily fully aligns with the direction of the fitness optimum (Fig. 4E).

**Figure 4.**
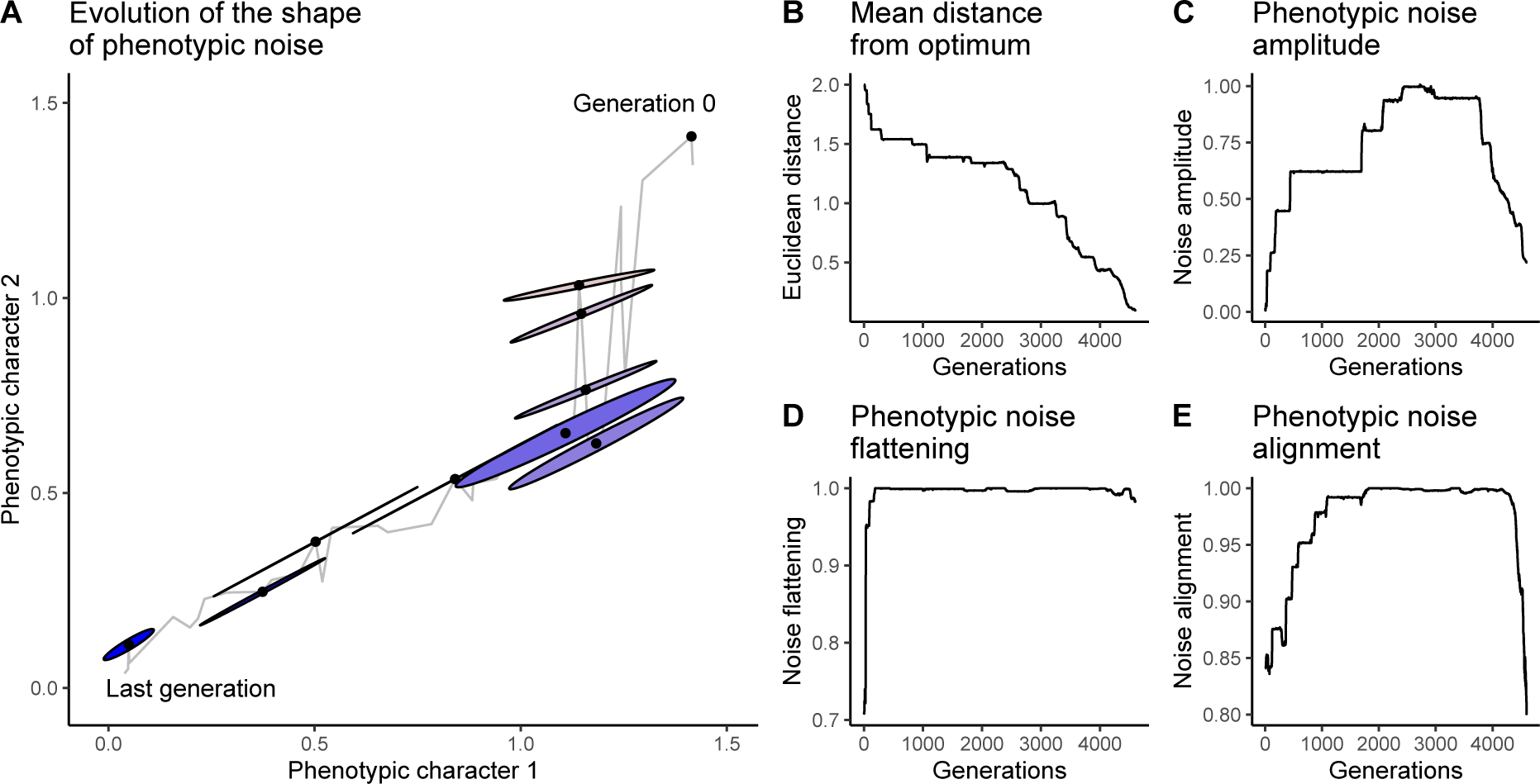
Simulation for two phenotypic characters under selection. In this simulation, a population of organisms with two phenotypic characters under selection evolves towards a fitness optimum ***z***_*opt*_ = (0, 0). The population size is *L* = 1, 000. Initial conditions are 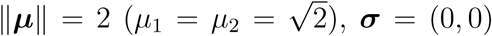 and ***θ*** = (0). Mutation sizes are *s*_*µ*_ = *s*_*σ*_ = *s*_*θ*_ = 0.1, mutation rates are *m*_*µ*_ = *m*_*σ*_ = *m*_*θ*_ = 10^−3^. The fitness function *w*(***z***) is Gaussian-shaped (*α* = 3.125, *β* = 0, *Q* = 2). **A**. Evolution of the phenotypic noise shape. Each ellipse represents noise amplitudes *σ*_1_ and *σ*_2_ (rescaled by a factor 0.3 for readability) and the orientation of the noise (given by ***θ***). The mean phenotype ***µ*** is represented by the central black dot. Generation time is represented by the color gradient, from beige to blue. The phenotypic noise shape of the best individual is plotted every 500 generations. The trajectory of the best individual of each generation is represented by a grey line. **B**. Distance of the population’s mean phenotype from the fitness optimum (*d* = ‖*µ‖*). **C**. Evolution of the population’s mean phenotypic noise amplitude (max(***σ***)). **D**. Evolution of the population’s mean phenotypic noise flattening. **E**. Evolution of the population’s mean phenotypic noise alignment with the direction of the fitness optimum.

### Phenotypic noise reduces the cost of complexity on the mean phenotype

Evolving an optimized phenotypic noise, fully correlated and aligned with the direction of the fitness optimum, is complex and requires to fix many mutations. As the cost of complexity impedes the production and the fixation of beneficial mutations for the mean phenotype, one could expect that this cost of complexity will also impede the probability to fix mutations shaping phenotypic noise in the right configuration. This probability depends at least on the phenotypic complexity *n*, the distance from the fitness optimum, the effective population size *N*_*e*_, and the genetic variability of ***µ, σ*** and ***θ***. When *β >* 0, if phenotypic noise evolves towards a single dimension aligned with the direction of the fitness optimum, it will increase the probability to fix beneficial mutations. This could counteract the cost of complexity on the mean phenotype and promote adaptive evolution.

To address the issue of the cost of complexity in the evolution of phenotypic noise, we simulated the adaptive evolution of initially maladapted populations towards the fitness optimum for a range of phenotypic complexities. We measured the convergence time to the fitness optimum in two fitness function configurations,

1. Gaussian-shaped fitness function (*α* = 3.125, *β* = 0 and *Q* = 2),
2. Rescaled minimum fitness function (*α* = 3.125, *β* = 0.1 and *Q* = 2).

We then varied the mutation rate ratio between the mean phenotype ***µ*** and the phenotypic noise parameters ***σ*** and ***θ*** (see Methods). We applied four scenarios:

1. Organisms have **no phenotypic noise**. Phenotypic noise mutation rates are *m*_*σ*_ = *m*_*θ*_ = 0. Phenotypic noise parameters are ***σ*** = **0, *θ*** = **0**. Mean phenotype mutation rate is *m*_*µ*_ = 10^−3^,
2. The phenotypic noise mutates **more slowly** than the mean phenotype. Phenotypic noise mutation rates are *m*_*σ*_ = *m*_*θ*_ = 10^−4^. Mean phenotype mutation rate is *m*_*µ*_ = 10^−3^,
3. The phenotypic noise and the mean phenotype mutate at the **same rate**. Mutation rates are *m*_*µ*_ = *m*_*σ*_ = *m*_*θ*_ = 10^−3^,
4. The phenotypic noise mutates **more often** than the mean phenotype. Phenotypic noise mutation rates are *m*_*σ*_ = *m*_*θ*_ = 10^−2^. Mean phenotype mutation rate is *m*_*µ*_ = 10^−3^.

The simulations were computed for a phenotypic complexity ranging from *n* = 1 to *n* = 10. 100 repetitions have been computed per parameter set (8,000 simulations in all). To limit computation time, the population was considered to have converged towards the fitness optimum when the mean phenotype in the population reached within one standard-deviation of the fitness optimum, *i.e*.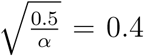. All the populations were initialized with a null phenotypic noise (***σ*** ∼ **0**), and the rotations of the covariance matrix set to ***θ*** = **0**. The initial Euclidean distance was ‖***µ***‖ = 2.0 (beyond the fitness landscape inflection point). Mutation sizes of phenotypic noise parameters and the mean phenotype were 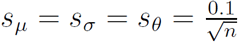 (the term 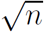 is used to keep a constant mutational size distribution whatever the phenotypic complexity). Simulation parameters are summarized in Table 4.

**Table 4.**
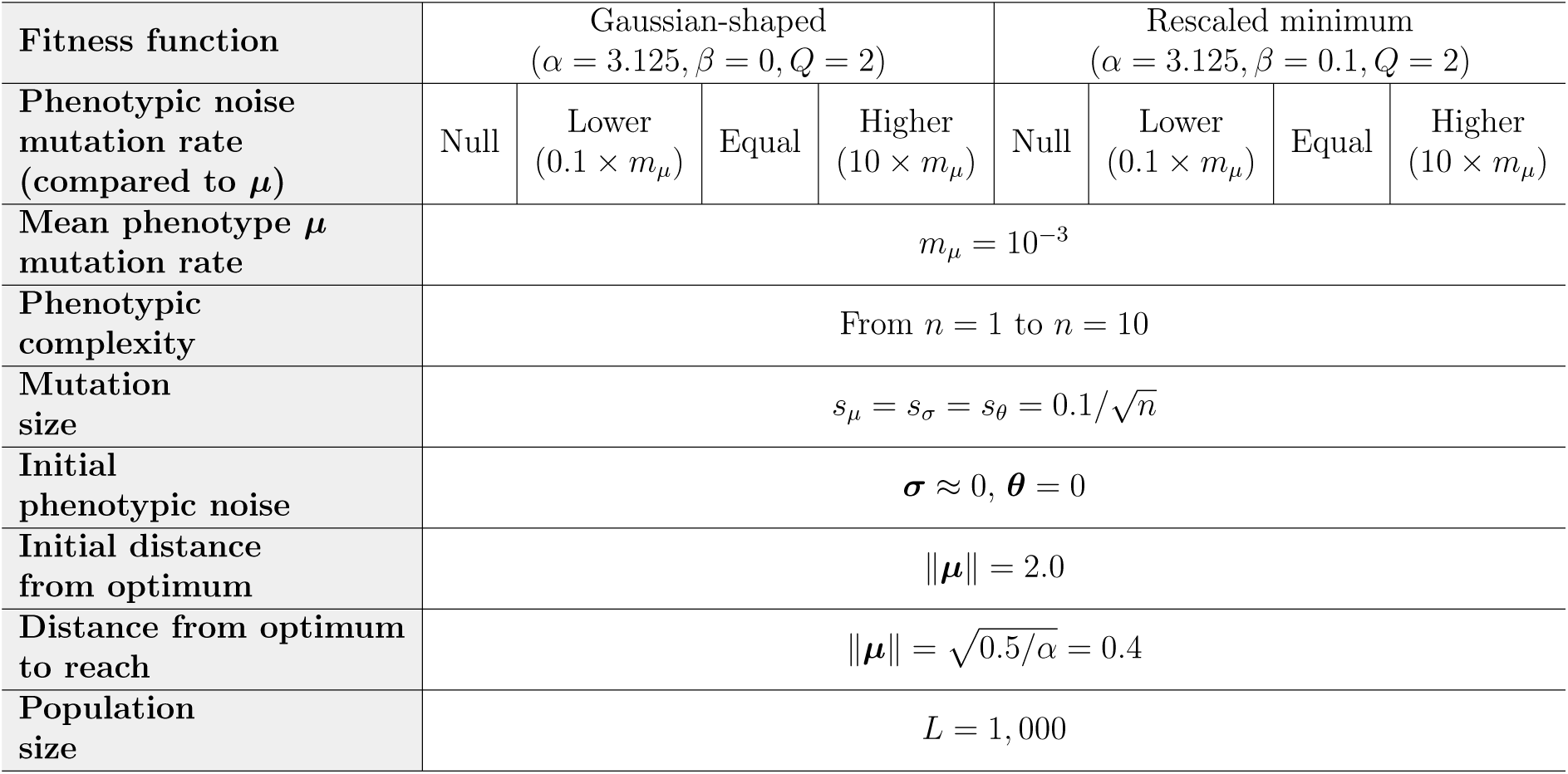
List of parameters used in the simulation framework for multiple phenotypic characters.

Figure 5 represents the convergence time of simulations, in number of generations (no phenotypic noise simulations: *brown curves*; lower mutation rate phenotypic noise simulations: *beige curves*; equal mutation rate phenotypic noise simulations: *water green curves*; higher mutation rate phenotypic noise simulations: *green curves*). While phenotypic noise systematically slows down adaptive evolution for a Gaussian-shaped fitness function (Fig. 5A), phenotypic noise significantly speeds up adaptive evolution for the rescaled minimum fitness function (Fig. 5B). The loss or gain of generation time depends on the mutation rate of phenotypic noise, the impact being stronger with a higher phenotypic noise mutation rate. **(i)** For the Gaussian-shaped fitness function (Fig. 5A), the convergence time of populations with the highest phenotypic noise mutation rate (Fig. 5A *green curve*) increases by more than 20 fold when *n* ≈ 3, although the difference with null phenotypic noise populations (Fig. 5A *brown curve*) fades away when the phenotypic complexity increases. The difference with null phenotypic noise populations is less significant for populations with a phenotypic noise mutation rate lower or equal to the mean phenotype mutation rate (Fig. 5A *beige and water green curves*). Around *n* ≈ 4, there is no more differences in convergence time between these populations. **(ii)** For the rescaled minimum fitness function (Fig. 5B), a very different trend is observed. The higher the phenotypic noise mutation rate is, the faster populations converge towards the fitness optimum. For populations with the highest phenotypic noise mutation rate (Fig. 5B *green curve*), there is a 10-fold decrease in convergence time compared to populations without phenotypic noise (Fig. 5B *brown curve*), indicating an important compensation of the cost of complexity. However, this gain is progressively lost when phenotypic complexity increases further.

**Figure 5.**
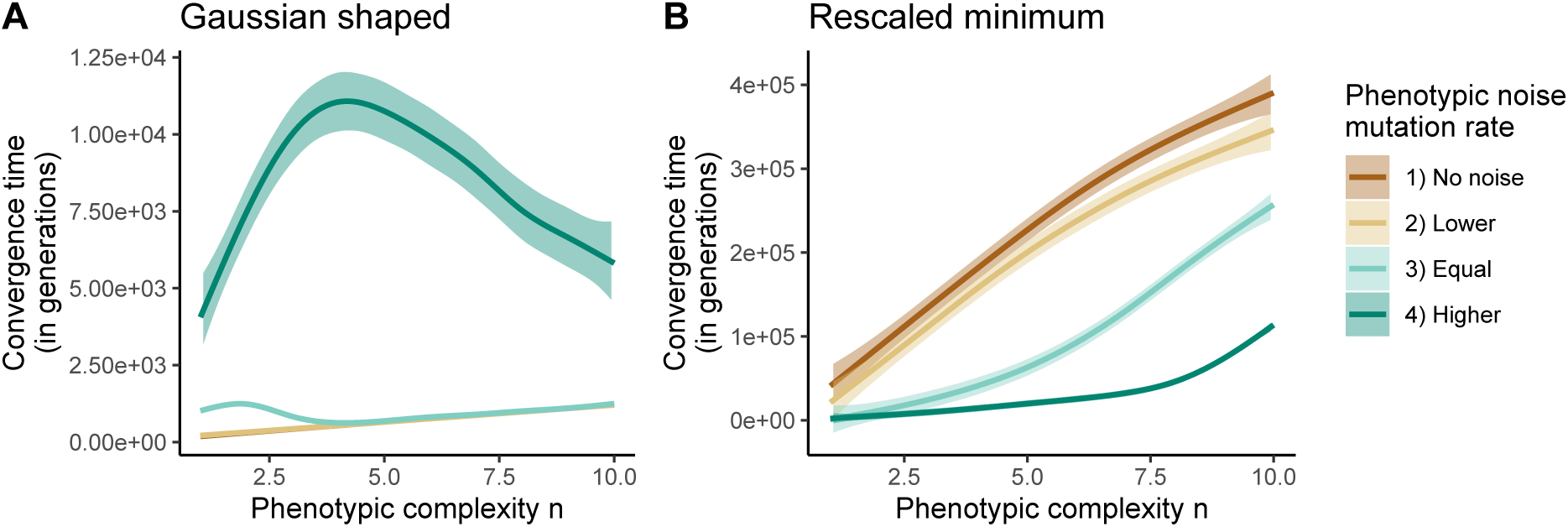
Convergence time of simulations in number of generations. **A**. Simulations with the Gaussian-shaped fitness function (*α* = 3.125, *β* = 0, *Q* = 2) **B**. Simulations with the rescaled minimum fitness function (*α* = 3.125, *β* = 0.1, *Q* = 2). The `lowess` smooth function from `R` is used to compute the mean and the standard error of the convergence time for each phenotypic complexity, with a span of 0.75. Brown line and standard error: populations with null phenotypic noise. Beige line and standard error: phenotypic noise mutation rate is lower than mean phenotype mutation rate. Water green line and standard error: phenotypic noise mutation rate is equal to the mean phenotype mutation rate. Green line and standard error: phenotypic noise mutation rate is higher than mean phenotype mutation rate. Due to computational limitations, the maximum number of generations was kept at 500,000.

To better characterize the evolution of phenotypic noise, we extracted for each simulation the maximal mean value reached in the population for the three indicators presented in Methods (phenotypic noise amplitude, flattening and alignment with the fitness optimum; Fig. 6). For the Gaussian-shaped fitness function (Figs. 6A-B-C), phenotypic noise gets closer to the optimal configuration when the mutation rate is higher, as shown by stronger phenotypic noise flattening and alignment (*i.e*. closer to 1). The optimality of the phenotypic noise configuration decreases with the phenotypic complexity: Indeed, as it is harder to evolve the right noise configuration when *n* increases, the mean phenotype of populations converges towards the fitness optimum before phenotypic noise is able to reach the best noise configuration. Nonetheless, for the populations with the highest phenotypic noise mutation rate (*green curves*), there is enough time for phenotypic noise to reach the optimal configuration if the phenotypic complexity is low (*n <* 4). In this case, phenotypic noise strongly reduces the gradient of relative genotypic fitness, increasing the convergence time (Fig. 5A).

**Figure 6.**
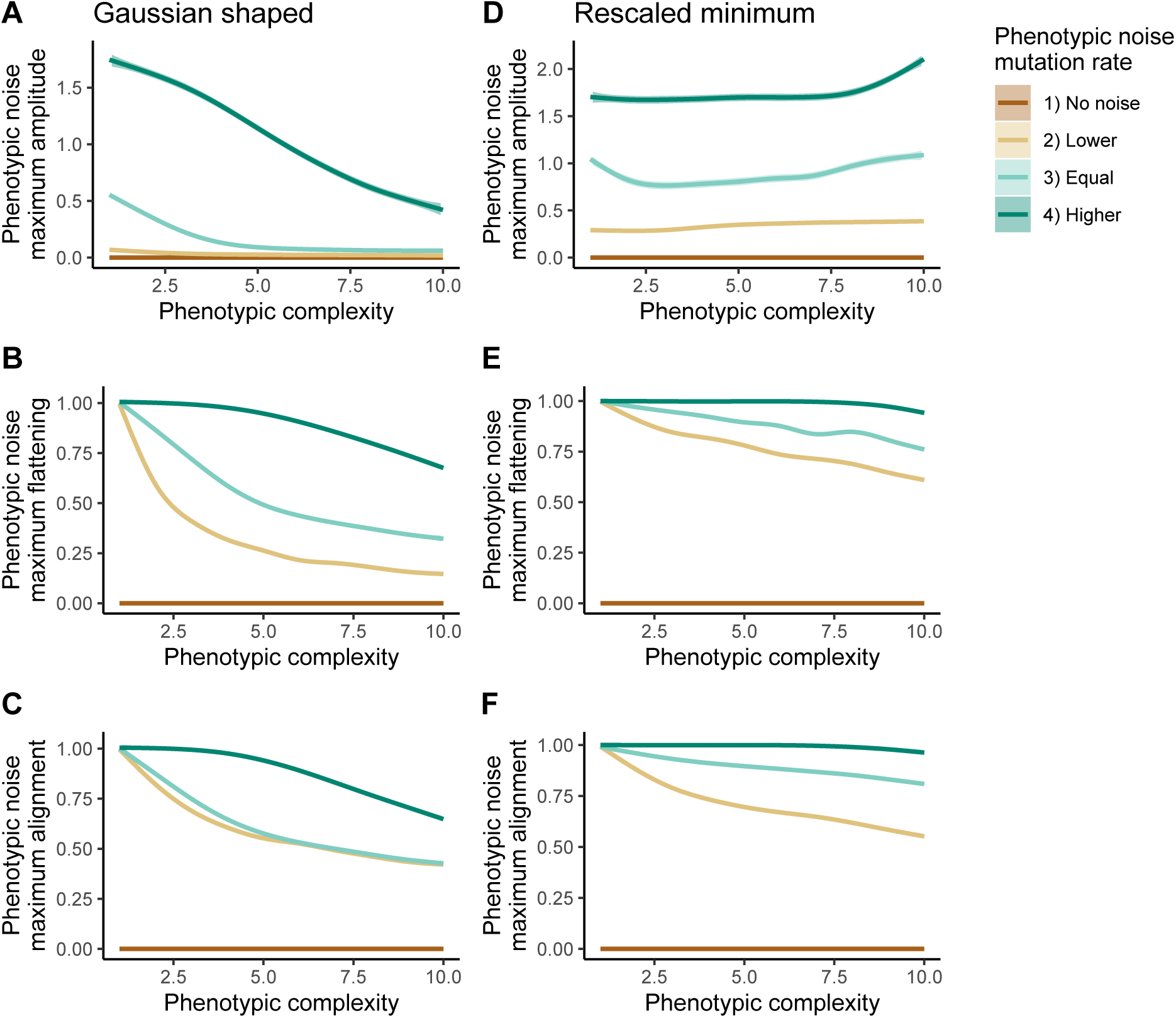
Mean optimal phenotypic noise configuration reached in the population for each scenario and each phenotypic complexity. **A**. Maximal phenotypic noise amplitude reached during simulations in Gaussian-shaped scenarios. **B**. Maximal phenotypic noise flattening reached during simulations in Gaussian-shaped scenarios. **C**. Maximal phenotypic noise alignment reached during simulations in Gaussian-shaped scenarios. **D**. Maximal phenotypic noise amplitude reached during simulations in rescaled minimum scenarios. **E**. Maximal phenotypic noise flattening reached during simulations in rescaled minimum scenarios. **F**. Maximal phenotypic noise alignment reached during simulations in rescaled minimum scenarios. Simulations are run for a phenotypic complexity ranging from *n* = 1 to *n* = 10. The `lowess` smooth function from `R` is used to compute the mean and the standard error of each measure for each phenotypic complexity, with a span of 0.75. Brown lines and standard errors: populations with null phenotypic noise. Beige lines and standard errors: phenotypic noise mutation rate is lower than mean phenotype mutation rate. Water green lines and standard errors: phenotypic noise mutation rate is equal to the mean phenotype mutation rate. Green lines and standard errors: phenotypic noise mutation rate is higher than mean phenotype mutation rate.

For the rescaled minimum fitness function (Figs. 6D-E-F), as the selection gradient is very low far from the optimum, phenotypic noise has enough time to evolve towards the optimal configuration, even for elevated phenotypic complexity. For populations with the highest phenotypic noise mutation rate (*green curves*), phenotypic noise always reaches the optimal configuration (noise flattening and alignment close to 1). These populations are by far the fastest to converge towards the fitness optimum (Fig. 5B), with a convergence time reduced by 90% compared to populations with no phenotypic noise. However, the cost of complexity increases on phenotypic noise evolution for high phenotypic complexity (*n >* 7). The lower the phenotypic noise mutation rate, the stronger the cost of complexity, as shown by weakened noise flattening and alignment when *n* increases. One can also notice that for high phenotypic complexity (*n >* 7), the phenotypic noise maximal amplitude increases. This can be explained by the time spent drifting away from the fitness optimum on the logarithmic fitness plateau. To summarize, phenotypic noise reduces the cost of complexity on the mean phenotype and promotes adaptive evolution as long as it does not suffer too much from the cost of complexity itself, and can reach the optimal configuration, aligned and fully correlated with the direction of the fitness optimum. According to our results, under the universal pleiotropy assumption, this is the case for an intermediate phenotypic complexity (*n* ranging from 1 to 7). For higher complexities, phenotypic noise suffers a significant cost, but still confers a strong advantage on the convergence time of populations.

## Discussion

While phenotypic noise has long been assumed to be harmful for the fitness, its benefit during an adaptation event was foreseen by Lande (1980), and since then studied, or even re-discovered, by many authors (Pal, 1998; Kawecki, 2000; Paenke et al., 2007; Zhang et al., 2009; Bruijning et al., 2019). Eldar and Elowitz (2010) suggested a verbal model where “*one might expect increased phenotypic noise during periods of adaptation to new environments, followed by reduction in noise when selection becomes stabilizing* “. However, until recently (Bódi et al., 2017; Duveau et al., 2018), the lack of experimental results to support the theory led to a certain confusion about the impact of phenotypic noise on organismal fitness during adaptation events. Indeed, it is assumed that phenotypic noise promotes adaptive evolution because it increases the genotypic fitness (Zhang et al., 2009). However, as a Gaussian-shaped fitness function has been historically used for the ease of calculation (Robertson, 1956), it is generally held that phenotypic noise only hinders the selection gradient (Lande, 1975). As suggested by Paenke et al. (2007), even if phenotypic noise increases the genotypic fitness far from the fitness optimum, it does not necessarily increase the gradient of relative genotypic fitness, and hence the selection gradient. This actually depends on the shape of the fitness function.

Here, using a quantitative genetics framework, we show that phenotypic noise promotes adaptive evolution on plateaus of positive fitness. Indeed, on a positive fitness plateau (*i.e*. on a log-fitness plateau), the local selection gradient is negligible. In the absence of phenotypic noise, the population drifts. However, a positive phenotypic noise generates a gradient of selection by magnifying the relative genotypic fitness differences present at the borders of the plateau, hence favoring the fixation of beneficial mutations.

This is a mechanism independent from the known effect of phenotypic noise on increasing the genotypic fitness and smoothing the absolute fitness landscape (Pál and Miklós, 1999; Saito et al., 2013). With our generalized unimodal fitness function (Eq. 1), this can lead to all sorts of scenarios depending on the shape of the fitness landscape (Table 2): **(i)** Far from the fitness optimum, phenotypic noise is always positively selected, and indirectly promotes adaptive evolution if the logarithmic fitness plateaus at a minimum of positive fitness (*β >* 0). One could object that such a fitness landscape is unrealistic, as there is no reason to not consider a null fitness very far from the optimum. However, in more complex fitness landscapes having multiple fitness optima, the existence of local plateaus or saddles, surrounded by valleys and peaks of fitness is likely. Regions of neutral evolution are proven phenomena in the evolution of DNA or protein sequences (reviewed by Payne and Wagner 2019). Phenotypic noise could then favor the fixation of rare beneficial mutations and phenotypic innovations, hence increasing evolvability (see *e.g*. Espinosa-Soto et al. 2011). **(ii)** Near the optimum, phenotypic noise is always selected against, while it has the potential to promote adaptive evolution when the logarithmic fitness plateaus at its maximum. This raises the question of the indirect selection of phenotypic noise for its specific impact on the gradient of relative genotypic fitness. Populations could fix mutations that increase the phenotypic noise by hitch hiking as a result of an enhanced rate of evolution. This could impact *e.g*. the evolution of bet-hedging strategies (when the location of the fitness optimum fluctuates through time, Beaumont et al. 2009; Bruijning et al. 2019), or more complex evolutionary outcomes such as the “survival of the flattest”, where a population undergoing a high mutation rate preferably evolves towards optima of intermediate fitness, but with a plateau reducing the fraction of deleterious mutations (Wilke et al., 2001). Although our study mainly provides theoretical arguments, recent experimental studies support our results. Bódi et al. (2017) developed inducible synthetic gene circuits to generate varying degrees of expression stochasticity of an anti-fungal resistance gene in *Saccharomyces cerevisiae*. They showed that phenotypic noise enhances the adaptive value of beneficial mutations when the expression stochasticity is higher. By altering the expression noise of the TDH3 gene in *Saccharomyces cerevisiae*, Duveau et al. (2018) showed that an increase in expression noise can be deleterious or beneficial, depending on the difference between the average expression level of a genotype and the expression level maximizing fitness: Far from the optimal expression level, expression noise is beneficial.

When organisms exhibit multiple phenotypic characters under stabilizing selection, the cost of complexity (Fisher, 1930; Orr, 2000) is expected to strongly reduce the fraction of beneficial mutations, and to slow down the rate of adaptive evolution towards the fitness optimum. Similarly, the impact of phenotypic noise on organismal fitness cannot simply be extrapolated from the single phenotypic character scenario. Our results show that when assuming a fully evolvable phenotypic noise in multiple dimensions (each dimension corresponding to a phenotypic character), the best phenotypic noise configuration far from the fitness optimum is aligned and fully correlated with the direction of the optimum, *i.e*. phenotypic noise undergoes a dimensionality reduction towards a single dimension. These results echo others showing that additive genetic correlations (the “G matrix”, Lande 1980) sometimes align towards the lines of least-resistance and the direction of selection (Penna et al., 2017), as suggested by Schluter (1996). A similar result exists for the evolution of the reaction-norm (Draghi and Whitlock, 2012; Lind et al., 2015; Gibert et al., 2019). Moreover, using numerical simulations, we show that when evolving populations reach such a phenotypic noise configuration, the fitness benefit of noise in one dimension is recovered. However, when the number of phenotypic characters under selection is too high (typically for *n >* 7), phenotypic noise suffer from complexity as well. While to our knowledge, no experimental study has shown that phenotypic noise promotes adaptive evolution for complex phenotypes, several authors have reported multi-characters phenotypic noise aligned with the direction of the selection in different species, such as *Drosophila serrata* (Sztepanacz et al., 2017) or *Daphnia* (Cressler et al., 2017). For example, Cressler et al. (2017) demonstrated the existence of correlated phenotypic noise on *Daphnia pulicaria* (a freshwater zooplankton). By measuring three integrated phenotypic characters at the individual level (body growth, number of eggs and longevity) on different populations of genetic variants, they showed that there was no significant genetic correlations between characters, while there is strong evidence for positive non-genetic correlations between characters: Increasing phenotypic noise enhances growth rate when non-genetic correlations between characters are positive, in agreement with our prediction on the evolution of phenotypic noise for multiple phenotypic characters. Experimental studies also suggest that both the amplitude and the correlation of phenotypic noise on multiple characters is evolvable (Stewart-Ornstein et al., 2012; Yvert et al., 2013). Moreover, the ability of the genotype-to-phenotype map to shape the random distribution of phenotypes, independently from the genetic variability, has been recently suggested by Sato and Kaneko (2019); Sakata and Kaneko (2020). They have shown with analytical and numerical approaches that thanks to the evolution of the genotype-to-phenotype map, the distribution of phenotypes in a population can be restricted to a subspace as a result of evolution in variables environments. All these results suggest that the evolution of a flattened phenotypic noise aligned with the fitness optimum is possible. Metzger et al. (2015) have also shown that sequences associated to expression noise in the gene TDH3 of *Saccharomyces cerevisiae* evolve faster that sequences associated to the mean expression level, suggesting that phenotypic noise could indeed evolve faster than the mean phenotype for some phenotypic characters.

Phenotypic noise is also a concern in cancer evolution, where it is known to facilitate the emergence of cancer resistance (Frank and Rosner, 2012; Huang, 2012; Pisco et al., 2013; Shaffer et al., 2017). Phenotypic noise could also play a role in the process of cell differentiation (Pujadas and Feinberg, 2012). Richard et al. (2016) showed that during the differentiation of chicken erythrocytes, gene expression patterns temporarily increase in variability, and then decrease when the cells reach the differentiated state. Although evolutionary arguments are delicate to apply to cell differentiation, one could speculate that evolution has shaped the epigenetic landscape such that differentiated phenotypic states are hills surrounded by flat regions where phenotypic noise could speed up cell differentiation.

Our study can be extended to more complex fitness landscapes, where the fitness value does not only depend on the distance from a single fitness optimum. Multi-peak and high-dimensional fitness landscapes containing plateaus of positive fitness could be used to evaluate the impact of an evolvable phenotypic noise on phenotypic innovation. Similarly, it would be interesting to study the impact of an evolvable phenotypic noise in variable environments having regions of plateauing fitness. It is also important to couple an evolvable phenotypic noise with evolvable genetic variability, mutation rates, and genetic regulation. Indeed, experimental results demonstrate the lack of realism of the universal pleiotropy assumption (see *e.g*. Wagner et al. 2008), and suggest that genetic variability could evolve towards a low dimensional distribution, just like phenotypic noise (Penna et al., 2017). Mutation rates are also known to evolve, at least temporarily, during adaptation events, as shown by the existence of mutator strains in micro-organisms (Taddei et al., 1997). Genetic regulation plays an important role in adaptive evolution as well, through the evolution of evolvability and robustness (see *e.g*. Crombach and Hogeweg 2008). The evolutive interaction of these multiple factors of genetic and phenotypic variability could lead to non-intuitive results.

Finally, our findings on the evolution of phenotypic noise during adaptation events could be used to predict the future direction of evolution, and to localize the direction of the fitness optimum in the phenotypic space. By tracking the evolution of phenotypic noise experimentally, it could be possible to find which selective pressures are at work on organisms, and to anticipate the next evolution steps. By deciphering the conditions in which phenotypic noise evolves towards specific patterns, our results may contribute to the growing field of predictive evolution.

## Supporting Information

**Figure S1. Shape of the fitness function** *w*(*z*) **depending on** *α, β* **and** *Q*.

**Figure S2. Simulated trajectories of populations with evolvable phenotypic noise**.

**Appendix S1. Analytical study of the impact of and evolvable phenotypic noise on fitness**.

**Script S1. Numerical approach to compute the isoclines** 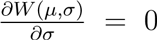 **and** 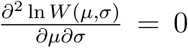.

**Script S2. Simulation framework**.

## References

Anderson, T. W., Olkin, I., and Underhill, L. G. (1987). Generation of random orthogonal matrices. SIAM Journal on Scientific and Statistical Computing, 8(4):625–629.

Beaumont, H. J., Gallie, J., Kost, C., Ferguson, G. C., and Rainey, P. B. (2009). Experimental evolution of bet hedging. Nature, 462(7269):90–93.

Bódi, Z., Farkas, Z., Nevozhay, D., Kalapis, D., Lázár, V., Csörgő, B., Nyerges, Á., Szamecz, B., Fekete, G., Papp, B., Araújo, H., Oliveira, J. L., Moura, G., Santos, M. A., Székely, T., Balázsi, G., and Pál, C. (2017). Phenotypic heterogeneity promotes adaptive evolution. PLOS Biology, 15(5):e2000644.

Boukhibar, L. M. and Barkoulas, M. (2016). The developmental genetics of biological robustness. Annals of Botany, 117(5):699–707.

Bruijning, M., Metcalf, C. J. E., Jongejans, E., and Ayroles, J. F. (2019). The evolution of variance control. Trends in ecology & evolution.

Cressler, C. E., Bengtson, S., and Nelson, W. A. (2017). Unexpected nongenetic individual heterogeneity and trait covariance in Daphnia And its consequences for ecological and evolutionary dynamics. The American Naturalist, 190(1):E13–E27.

Crombach, A. and Hogeweg, P. (2008). Evolution of evolvability in gene regulatory networks. PLoS computational biology, 4(7).

Draghi, J. (2019). Phenotypic variability can promote the evolution of adaptive plasticity by reducing the stringency of natural selection. Journal of evolutionary biology.

Draghi, J. A. and Whitlock, M. C. (2012). Phenotypic plasticity facilitates mutational variance, genetic variance, and evolvability along the major axis of environmental variation. Evolution: International Journal of Organic Evolution, 66(9):2891–2902.

Duveau, F., Hodgins-Davis, A., Metzger, B. P., Yang, B., Tryban, S., Walker, E. A., Lybrook, T., and Wittkopp, P. J. (2018). Fitness effects of altering gene expression noise in saccharomyces cerevisiae. Elife, 7:e37272.

Eldar, A. and Elowitz, M. B. (2010). Functional roles for noise in genetic circuits. Nature, 467(7312):167–173.

Eling, N., Morgan, M. D., and Marioni, J. C. (2019). Challenges in measuring and understanding biological noise. Nature Reviews Genetics, page 1.

Elowitz, M. B., Levine, A. J., Siggia, E. D., and Swain, P. S. (2002). Stochastic Gene Expression in a Single Cell. Science, 297(5584):1183–1186.

Espinosa-Soto, C., Martin, O. C., and Wagner, A. (2011). Phenotypic plasticity can facilitate adaptive evolution in gene regulatory circuits. BMC evolutionary biology, 11(1):5.

Falconer, D. and Robertson, A. (1956). Selection for environmental variability of body size in mice. Molecular and General Genetics MGG, 87(3):385–391.

Fisher, R. A. (1930). The genetical theory of natural selection: a complete variorum edition. Oxford University Press, Oxford (UK).

Forsman, A. (2015). Rethinking phenotypic plasticity and its consequences for individuals, populations and species. Heredity, 115(4):276.

Frank, S. A. and Rosner, M. R. (2012). Nonheritable cellular variability accelerates the evolutionary processes of cancer. PLoS biology, 10(4):e1001296.

Fraser, H. B., Hirsh, A. E., Giaever, G., Kumm, J., and Eisen, M. B. (2004). Noise minimization in eukaryotic gene expression. PLoS Biology, 2(6):834–838.

Gavrilets, S. and Hastings, A. (1994). A quantitative-genetic model for selection on developmental noise. Evolution, 48(5):1478–1486.

Gibert, P., Debat, V., and Ghalambor, C. K. (2019). Phenotypic plasticity, global change, and the speed of adaptive evolution. Current opinion in insect science.

Hill, W. G. and Mulder, H. A. (2010). Genetic analysis of environmental variation. Genetics Research, 92(5-6):381–395.

Hortsch, S. K. and Kremling, A. (2018). Characterization of noise in multistable genetic circuits reveals ways to modulate heterogeneity. PloS one, 13(3):e0194779.

Huang, S. (2012). Tumor progression: Chance and necessity in Darwinian and Lamarckian somatic (mutationless) evolution. Progress in Biophysics and Molecular Biology, 110(1):69–86.

Ito, Y., Toyota, H., Kaneko, K., and Yomo, T. (2009). How selection affects phenotypic fluctuation. Molecular Systems Biology, 5(264):264.

Kawecki, T. J. (2000). The evolution of genetic canalization under fluctuating selection. Evolution, 54(1):1–12.

Keren, L., Hausser, J., Lotan-Pompan, M., Vainberg Slutskin, I., Alisar, H., Kaminski, S., Weinberger, A., Alon, U., Milo, R., and Segal, E. (2016). Massively Parallel Interrogation of the Effects of Gene Expression Levels on Fitness. Cell, 166(5):1282–1294.e18.

Lande, R. (1975). The maintenance of genetic variability by mutation in a polygenic character with linked loci. Genetics Research, 26(3):221–235.

Lande, R. (1976). Natural selection and random genetic drift in phenotypic evolution. Evolution, 30(2):314–334.

Lande, R. (1979). Quantitative genetic analysis of multivariate evolution, applied to brain: body size allometry. Evolution, 33(1Part2):402–416.

Lande, R. (1980). The genetic covariance between characters maintained by pleiotropic mutations. Genetics, 94(1):203–215.

Lehner, B. (2008). Selection to minimise noise in living systems and its implications for the evolution of gene expression. Molecular Systems Biology, 4(170):170.

Lind, M. I., Yarlett, K., Reger, J., Carter, M. J., and Beckerman, A. P. (2015). The alignment between phenotypic plasticity, the major axis of genetic variation and the response to selection. Proceedings of the Royal Society B: Biological Sciences, 282(1816):20151651.

Martin, G. (2014). Fisher’s geometrical model emerges as a property of complex integrated phenotypic networks. Genetics, 197(1):237–255.

Martin, G. and Lenormand, T. (2006). A general multivariate extension of Fisher’s geometrical model and the distribution of mutation fitness effects across species. Evolution, 60(5):893–907.

Metzger, B. P. H., Yuan, D. C., Gruber, J. D., Duveau, F. D., and Wittkopp, P. J. (2015). Selection on noise constrains variation in a eukaryotic promoter. Nature, 521(521):344–347.

Mineta, K., Matsumoto, T., Osada, N., and Araki, H. (2015). Population genetics of non-genetic traits: Evolutionary roles of stochasticity in gene expression. Gene, 562(1):16–21.

Mulder, H. A., Gienapp, P., and Visser, M. E. (2016). Genetic variation in variability: phenotypic variability of fledging weight and its evolution in a songbird population. Evolution, 70(9):2004–2016.

Newman, J. R. S., Ghaemmaghami, S., Ihmels, J., Breslow, D. K., Noble, M., DeRisi, J. L., and Weissman, J. S. (2006). Single-cell proteomic analysis of S. cerevisiae reveals the architecture of biological noise. Nature, 441(7095):840–846.

Ohya, Y., Kimori, Y., Okada, H., and Ohnuki, S. (2015). Single-cell phenomics in budding yeast. Molecular Biology of the Cell, 26(22):3920–3925.

Ordas, B., Malvar, R. A., and Hill, W. G. (2008). Genetic variation and quantitative trait loci associated with developmental stability and the environmental correlation between traits in maize. Genetics research, 90(5):385–395.

Orr, H. A. (2000). Adaptation and the Cost of Complexity. Evolution, 54(1):13–20.

Orr, H. A. (2005). The genetic theory of adaptation: a brief history. Nature Reviews Genetics, 6(2):119.

Paenke, I., Sendhoff, B., and Kawecki, T. J. (2007). Influence of plasticity and learning on evolution under directional selection. The American Naturalist, 170(2):E47–E58.

Pal, C. (1998). Plasticity, memory and the adaptive landscape of the genotype. Proceedings of the Royal Society of London. Series B: Biological Sciences, 265(1403):1319–1323.

Pál, C. and Miklós, I. (1999). Epigenetic inheritance, genetic assimilation and speciation. Journal of Theoretical Biology, 200(1):19–37.

Payne, J. L. and Wagner, A. (2019). The causes of evolvability and their evolution. Nature Reviews Genetics, 20(1):24–38.

Pelabon, C., Hansen, T. F., Carter, A. J., and Houle, D. (2010). Evolution of variation and variability under fluctuating, stabilizing, and disruptive selection. Evolution: International Journal of Organic Evolution, 64(7):1912–1925.

Penna, A., Melo, D., Bernardi, S., Oyarzabal, M. I., and Marroig, G. (2017). The evolution of phenotypic integration: How directional selection reshapes covariation in mice. Evolution, 71(10):2370–2380.

Pisco, A. O., Brock, A., Zhou, J., Moor, A., Mojtahedi, M., Jackson, D., and Huang, S. (2013). Non-Darwinian dynamics in therapy-induced cancer drug resistance. Nature Communications, 4:2467.

Pujadas, E. and Feinberg, A. P. (2012). Regulated noise in the epigenetic landscape of development and disease. Cell, 148(6):1123–1131.

Raser, J. M. and O’Shea, E. K. (2005). Noise in Gene Expression: Orgins, Consequences, and Control. Science, 309(5743):2010–2013.

Reyes, J., Chen, J.-Y., Stewart-Ornstein, J., Karhohs, K. W., Mock, C. S., and Lahav, G. (2018). Fluctuations in p53 signaling allow escape from cell-cycle arrest. Molecular cell, 71(4):581–591.

Richard, A., Boullu, L., Herbach, U., Bonnafoux, A., Morin, V., Vallin, E., Guillemin, A., Gao, N. P., Gunawan, R., Cosette, J., et al. (2016). Single-cell-based analysis highlights a surge in cell-to-cell molecular variability preceding irreversible commitment in a differentiation process. PLoS biology, 14(12):e1002585.

Robertson, A. (1956). The effect of selection against extreme deviants based on deviation or on homozygosis. Journal of Genetics, 54(2):236.

Saito, N., Ishihara, S., and Kaneko, K. (2013). Baldwin effect under multipeaked fitness landscapes: phenotypic fluctuation accelerates evolutionary rate. Physical Review E, 87(5):052701.

Sakata, A. and Kaneko, K. (2020). Dimensional reduction in evolving spin-glass model: correlation of phenotypic responses to environmental and mutational changes. arXiv preprint 2001.03714.

Sato, T. U. and Kaneko, K. (2019). Evolutionary dimension reduction in phenotypic space. arXiv preprint 1910.01297.

Schluter, D. (1996). Adaptive radiation along genetic lines of least resistance. Evolution, 50(5):1766–1774.

Schmutzer, M. and Wagner, A. (2020). Gene expression noise can promote the fixation of beneficial mutations in fluctuating environments. bioRxiv.

Shaffer, S. M., Dunagin, M. C., Torborg, S. R., Torre, E. A., Emert, B., Krepler, C., Beqiri, M., Sproesser, K., Brafford, P. A., Xiao, M., et al. (2017). Rare cell variability and drug-induced reprogramming as a mode of cancer drug resistance. Nature, 546(7658):431.

Shen, X., Pettersson, M., Rönnegård, L., and Carlborg, Ö. (2012). Inheritance beyond plain heritability: variance-controlling genes in arabidopsis thaliana. PLoS genetics, 8(8):e1002839.

Stewart-Ornstein, J., Weissman, J. S., and El-Samad, H. (2012). Cellular noise regulons underlie fluctuations in saccharomyces cerevisiae. Molecular cell, 45(4):483–493.

Sztepanacz, J. L., McGuigan, K., and Blows, M. W. (2017). Heritable micro-environmental variance covaries with fitness in an outbred population of drosophila serrata. Genetics, 206(4):2185–2198.

Taddei, F., Radman, M., Maynard-Smith, J., Toupance, B., Gouyon, P.-H., and Godelle, B. (1997). Role of mutator alleles in adaptive evolution. Nature, 387(6634):700–702.

Tenaillon, O. (2014). The utility of fisher’s geometric model in evolutionary genetics. Annual review of ecology, evolution, and systematics, 45:179–201.

Viñuelas, J., Kaneko, G., Coulon, A., Beslon, G., and Gandrillon, O. (2012). Towards experimental manipulation of stochasticity in gene expression. Progress in biophysics and molecular biology, 110(1):44–53.

Wagner, G. P., Kenney-Hunt, J. P., Pavlicev, M., Peck, J. R., Waxman, D., and Cheverud, J. M. (2008). Pleiotropic scaling of gene effects and the ‘cost of complexity’. Nature, 452(7186):470–472.

Wang, Z. and Zhang, J. (2011). Impact of gene expression noise on organismal fitness and the efficacy of natural selection. Proceedings of the National Academy of Sciences of the United States of America, 108(16):E67–76.

Welch, J. J. and Waxman, D. (2003). Modularity and the cost of complexity. Evolution, 57(8):1723–1734.

Wilke, C. O., Wang, J. L., Ofria, C., Lenski, R. E., and Adami, C. (2001). Evolution of digital organisms at high mutation rates leads to survival of the flattest. Nature, 412(6844):331–333.

Yvert, G., Ohnuki, S., Nogami, S., Imanaga, Y., Fehrmann, S., Schacherer, J., and Ohya, Y. (2013). Single-cell phenomics reveals intra-species variation of phenotypic noise in yeast. BMC Systems Biology, 7(1):54.

Zhang, Z., Qian, W., and Zhang, J. (2009). Positive selection for elevated gene expression noise in yeast. Molecular Systems Biology, 5(1):299.

