## Supplementary material for "Phenotypic Noise and the Cost of Complexity": Figure S1

### Phenotypic Noise and the Cost of Complexity.
### FIGURE S1—Shape of the fitness function $w(z)$ depending on $\alpha$, $\beta$ and $Q$

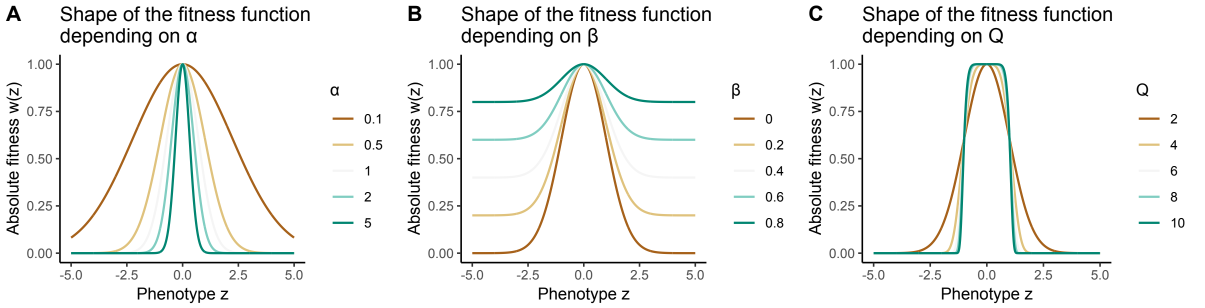


Figure S1 – **Shape of the fitness function** $w(z)$ **depending on** $\alpha$**,** $\beta$ **and** $Q$**.** The shape of the generalized fitness function $w(z) = (1-\beta)e^{-\alpha z^Q} + \beta$ depends on the parameters $\alpha$, $\beta$ and $Q$. **A.** Shape of $w(z)$ depending on $\alpha$. **B.** Shape of $w(z)$ depending on $\beta$. **C.** Shape of $w(z)$ depending on $Q$. Default parameters are $\alpha = 0.5$, $\beta = 0.1$ and $Q = 2$.
