## Supplementary material for "Phenotypic Noise and the Cost of Complexity": Figure S2

### Phenotypic Noise and the Cost of Complexity. FIGURE S2—Simulated trajectories of populations with evolvable phenotypic noise

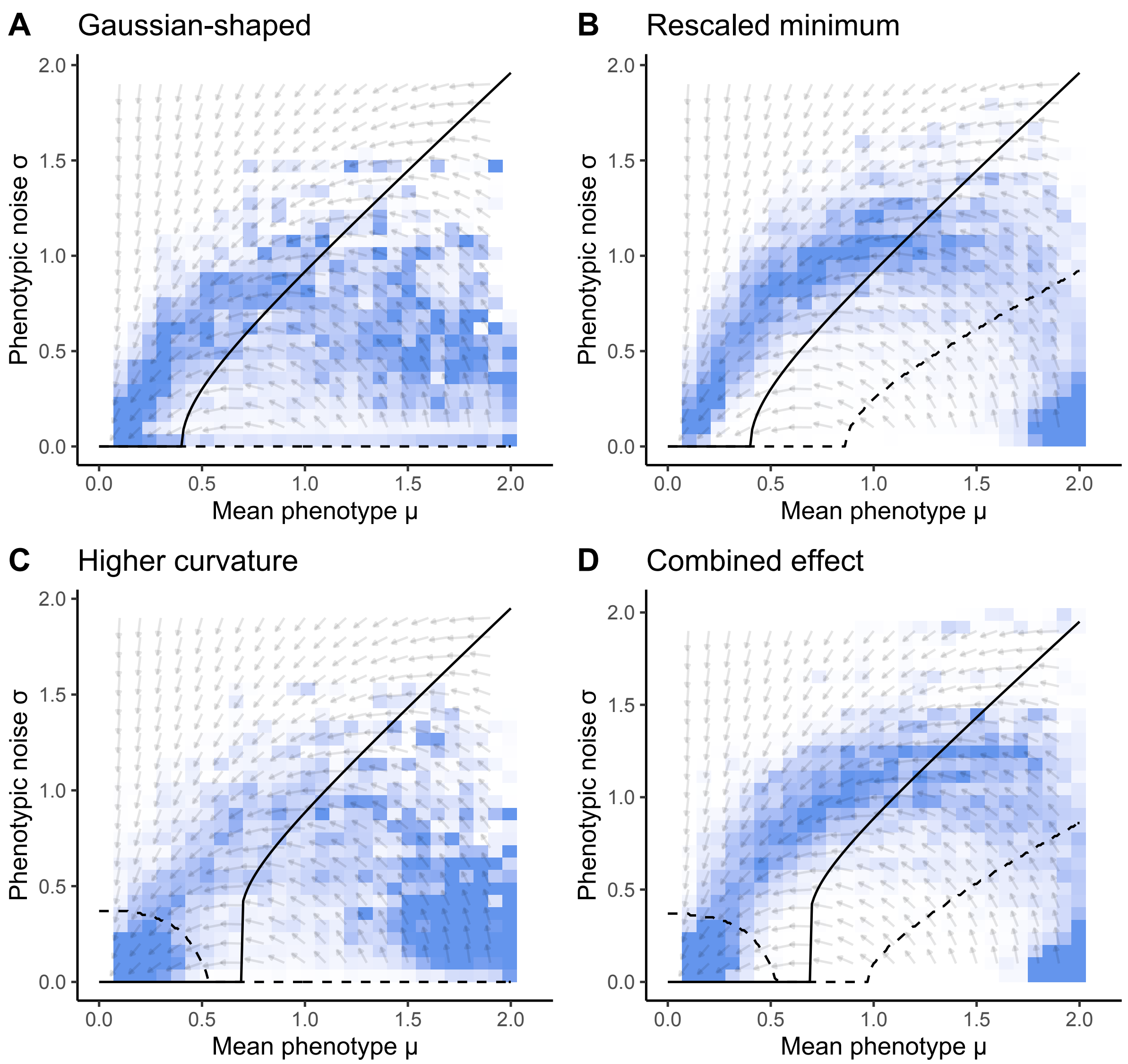


Figure S2 – **Simulated trajectories of populations with evolvable phenotypic noise.** The shape of the generalized fitness function $w(z) = (1-\beta)e^{-\alpha z^Q} + \beta$ depends on the parameters $\alpha$, $\beta$ and $Q$. **A.** Gaussian-shaped fitness landscape ($\alpha = 3.125$, $\beta = 0$ and $Q = 2$). **B.** Rescaled minimum fitness landscape ($\alpha = 3.125$, $\beta = 0.1$ and $Q = 2$). **C.** Higher curvature fitness landscape ($\alpha = 3.125$, $\beta = 0$ and $Q = 4$). **D.** Combined effects fitness landscape ($\alpha = 3.125$, $\beta = 0.1$ and $Q = 4$). For each scenario, the 200 repetitions are summarized by a density heatmap. The isocline $dW/d\sigma = 0$ for which $\sigma$ maximizes $W(\mu, \sigma)$ is represented by a solid black line. The isocline $\frac{\partial \ln W(\mu,\sigma)}{\partial\mu\partial\sigma} = 0$ for which $\sigma$ maximizes $\frac{\partial \ln W(\mu,\sigma)}{\partial\mu}$ is represented by a dashed black line. The direction of the gradient of relative genotypic fitness in the space $(\mu, \sigma)$ is represented by a vector field.
