## Appendix S1 for "Phenotypic Noise and the Cost of Complexity"

### Phenotypic Noise and the Cost of Complexity. Appendix S1—Analytical study of the impact of and evolvable phenotypic noise on fitness

#### Contents


**1 Inflection point of the fitness function $w(z)$, depending on parameters $\alpha$ and $Q$ 2**

**2 Impact of phenotypic noise on the gradient of relative genotypic fitness 4**

**3 For multiple characters under selection, the best phenotypic noise configuration is fully correlated and aligned with the fitness optimum 10**


#### List of Figures


1 Phenotypic noise increases the gradient of relative genotypic fitness when the logarithmic fitness plateaus. . . . . . . . . . . . . . . . . . . . . . . . . . 9

2 Anisotropic and correlated phenotypic noise for two phenotypic characters under selection. . . . . . . . . . . . . . . . . . . . . . . . . . . . . . . . . 13

### 1 Inflection point of the fitness function $w(z)$, depending on parameters $\alpha$ and $Q$

We consider a large population of individuals under stabilizing selection and with discrete generations. We define each individual's genotype by the pair $\{\mu, \sigma\}$, $\mu \in \mathbb{R}$ being the mean phenotype (or breeding value) and $\sigma \in \mathbb{R}^+$ the phenotypic noise amplitude. Both $\mu$ and $\sigma$ are controlled genetically, and evolvable. Individual continuous phenotype $z \in \mathbb{R}$ follows a normal distribution of mean $\mu$ and variance $\sigma^2$: $p(z|\mu, \sigma^2) \sim \mathcal{N}(\mu, \sigma^2)$. Individual absolute fitness $w$ depends on the phenotype $z$. The fitness function $w(z)$ is assumed positive, symmetric, three times differentiable and possessing at least one non-degenerate optimum (Martin, 2014). We chose $w$ as the unimodal function

$$w(z) = (1 - \beta)e^{-\alpha z^Q} + \beta. \tag{1}$$

$\alpha > 0$ controls the sharpness of the fitness peak at $z = 0$. $\beta \in [0, 1[$ controls the minimal fitness, when $|z| \gg 0$. The curvature parameter $Q = 2k$, for positive integers $k$, controls how close the fitness peak is from a step function (when $Q = +\infty$). When $\beta = 0$ and $Q = 2$, $w$ is a Gaussian fitness function with variance $1/(2\alpha)$. The absolute fitness of a genotype $\{\mu, \sigma\}$ (or genotypic fitness) is the expected fitness across all possible phenotypes given the phenotypic noise amplitude $\sigma$ (Lande, 1979),

$$W(\mu, \sigma) = \int_{-\infty}^{+\infty} p(z|\mu, \sigma^2) w(z) dz. \tag{2}$$

Using a Taylor expansion, it has been shown (Paenke et al., 2007) that the genotypic fitness (Eq. 2) can be expressed by:

$$W(\mu, \sigma) = w(\mu) + \frac{\sigma^2}{2} w''(\mu) + O(\sigma^4), \tag{3}$$

where $w''$ is the second derivative of $w$. Hence, the effect of small phenotypic noise (such that $O(\sigma^4)$ remains negligible) on genotypic fitness is determined by the local curvature: when $w(\mu)$ is convex ($w''(\mu) > 0$), phenotypic noise increases the genotypic fitness. When $w(\mu)$ is concave ($w''(\mu) < 0$), phenotypic noise decreases the genotypic fitness. This is a well-known result (Slatkin and Lande, 1976), which can also be obtained directly from the definition of $W$ under weaker smoothness assumption, via Jensen's Inequality.

The first derivative of $w$ is

$$w'(z) = -\alpha Q z^{Q-1}(w - \beta). \tag{4}$$

The term $(w - \beta)$ is always positive, and $\operatorname{sign} w'(z) = \operatorname{sign} -z$, as expected. The second derivative of $w$ is

$$w''(z) = \alpha Q z^{Q-2}(w - \beta)\big[\alpha Q z^Q - (Q - 1)\big]. \tag{5}$$

The inflection points of $w$ are given by

$$z^Q = \frac{(Q - 1)}{\alpha Q}. \tag{6}$$

When $\alpha$ increases, the fitness peak gets sharper and inflection points move closer to the fitness optimum ($z = 0$). When the curvature $Q$ increases, the fitness function approaches a step function, and the inflection points converge to $z_{\text{infl}} = \pm 1$. The minimal fitness $\beta$ has no effect on the position of the inflection points.

In summary, **the phenotypic noise $\sigma$ increases the genotypic fitness $W(\mu, \sigma)$ when the mean phenotype $\mu$ is beyond the inflection point of $w(\mu)$**, which is given by Equation 6. However, this analysis does not provide the optimal value of $\sigma$ that maximizes $W(\mu, \sigma)$ (except below the inflection point, where the optimal value of $\sigma$ is zero). To do so, we must compute numerically $\sigma$ such that $\frac{\partial W(\mu,\sigma)}{\partial \sigma} = 0$ (see main manuscript).

### 2 Impact of phenotypic noise on the gradient of relative genotypic fitness

As discussed by Paenke et al. (2007), if all the genetic variation is additive or if reproduction is asexual, the response to selection equals the selection differential, *i.e.* the difference of the base population mean and the mean after selection (Lande, 1979). This does not hold with dominance or epistasis, but it is reasonable to assume that in most cases the response to selection will be positively correlated with the selection differential. Moreover, Paenke et al. (2007) have shown that if the effect of phenotypic noise on the genotypic fitness $W(\mu, \sigma)$ is monotonic, which holds when genotypic variability is small, it is sufficient to study the impact of phenotypic noise on the gradient of relative genotypic fitness $\frac{\partial \ln W(\mu,\sigma)}{\partial \mu}$.

Using a Taylor expansion around $\sigma = 0$, we show that the effect of phenotypic noise amplitude $\sigma$ on the gradient of relative genotypic fitness, *i.e.* $\frac{\partial^2 \ln W(\mu,\sigma)}{\partial \mu \partial \sigma}$, is given by

$$\frac{\partial^2 \ln W(\mu, \sigma)}{\partial \mu \partial \sigma} \approx \frac{w'(\mu)}{w(\mu)} + \frac{\sigma^2}{2w(\mu)^2} \left[w(\mu)w'''(\mu) - w'(\mu)w''(\mu)\right] + O(\sigma^3). \tag{7}$$

Thus, the phenotypic noise amplitude $\sigma$ can increase the gradient of relative genotypic fitness, and promotes adaptive evolution, provided that (Paenke et al., 2007)

$$\operatorname{sign}\big(w(\mu)w'''(\mu) - w'(\mu)w''(\mu)\big) = \operatorname{sign} w'(\mu). \tag{8}$$

Because the fitness function is assumed symmetric, we can restrict the study to $\mu \leq 0$. For simplicity, we further assume that the fitness function is increasing for $\mu < 0$. For $\mu < 0$, $w'(\mu) > 0$, and the condition for phenotypic noise to promote evolution is $[w(\mu)w'''(\mu) - w'(\mu)w''(\mu)] > 0$ (Eq. 8). Except on the inflection point where $w''(\mu) = 0$, this condition is equivalent to

$$\begin{cases} \dfrac{w'''}{w''} < \dfrac{w'}{w}, & \text{if } w'' < 0, \\ \\ \dfrac{w'''}{w''} > \dfrac{w'}{w}, & \text{if } w'' > 0. \end{cases} \tag{9}$$

This can be expressed in terms of log-derivatives

$$\begin{cases} \dfrac{d\ln(-w'')}{d\mu} < \dfrac{d\ln(w)}{d\mu}, & \text{if } w'' < 0, \\ \\ \dfrac{d\ln(w)}{d\mu} < \dfrac{d\ln(w'')}{d\mu}, & \text{if } w'' > 0, \end{cases} \tag{10}$$

(the minus sign is there to keep the argument of the log positive). $w'' < 0$ occurs close to the fitness optimum, while $w'' > 0$ occurs far from the optimum. The conditions are the same for $\mu > 0$, if the derivative is taken with respect to $-\mu$ (*i.e.* towards the fitness optimum). Therefore we have a general condition for phenotypic noise to increase the selection gradient. When moving towards the fitness optimum, phenotypic noise will promote adaptive evolution if,

— close to the fitness optimum (when $w'' < 0$), the relative curvature decreases, or increases more slowly than the relative fitness.

— far from the optimum (when $w'' > 0$), the relative curvature increases more rapidly than the relative fitness.

Below, we show that none of these conditions can be met with a Gaussian fitness function (*i.e.* when $\beta = 0$ and $Q = 2$ in Eq. 1): in this case, the relative change in fitness is proportional to $|\mu|$. Far from the optimum, it is too large compared to relative change in curvature. Close to the optimum, the relative curvature increases more rapidly than the relative fitness. However, with other kind of unimodal fitness functions, two cases allow phenotypic noise to promote adaptive evolution: **(i)** When the fitness function plateaus near the fitness optimum ($Q > 2$), the relative curvature decreases when approaching the optimum. This means that phenotypic noise will speed up evolution in a plateauing landscape around the optimum. **(ii)** When the fitness function is strictly bounded below by a positive value ($\beta > 0$), the relative change in fitness tends to zero far from the optimum, and it becomes possible for the relative curvature to increase faster than it. Finally, in the neighborhood of the inflection point $w'' \sim 0$, the condition 8 becomes just $w(\mu)w''' > 0$. However, we expect generically negative third derivative to compensate for the positive first derivative: indeed, around the inflection point, the fitness function behaves as a sigmoidal function.

Therefore, previous arguments imply that phenotypic noise will be detrimental to the gradient of relative genotypic fitness except in two cases:

**(i)** Around a flat fitness optimum,

**(ii)** Near a positive fitness minimum.

To predict regions of $w(z)$ where phenotypic noise promotes adaptive evolution, we need to make these statements more quantitative. To do so, one needs to compute the third derivative of $w$,

$$w'''(z) = \begin{cases} 4\alpha^2 z(w-\beta)\big[3-2\alpha z^2\big], & Q=2, \\ \alpha Q z^{Q-3}(w-\beta)\big[\alpha Q z^Q(3Q-3) & \\ -\alpha^2 Q^2 z^{2Q} - (Q-1)(Q-2)\big], & Q \geq 4. \end{cases} \tag{11}$$

- **First**, we treat the Gaussian case ($Q = 2, \beta = 0$).

$$w'''w - w'w'' = 8\alpha^2 \mu w^2. \tag{12}$$

The sign is always opposite to $w'$, so condition 8 can never be satisfied.

- **Second**, if $Q = 2, \beta > 0$,

$$w'''w - w'w'' = 4\alpha^2 \mu (w-\beta)\big[2w - 2\alpha\beta\mu^2 + \beta\big]. \tag{13}$$

Condition 8 will be satisfied if $\left[2w - 2\alpha\beta\mu^2 + \beta\right] < 0$. This occurs, and phenotypic noise promotes adaptive evolution if

$$\mu^2 > \frac{2w + \beta}{2\alpha\beta}. \tag{14}$$

- **Third**, if $Q > 2, \beta = 0$,

$$w'''w - w'w'' = \alpha Q \mu^{Q-3} w^2 \Big[2w(Q-1)\alpha Q\mu^Q - w(Q-1)(Q-2)\Big]. \tag{15}$$

Condition 8 will be satisfied if the term in the square brackets is negative, that is if

$$\mu^Q < \frac{Q-2}{2\alpha Q}. \tag{16}$$

- **Fourth**, if $Q > 2, \beta > 0$,

$$\begin{aligned} w'''w - w'w'' = \alpha Q\mu^{Q-3} w(w-\beta)\Big[&-\alpha^2\beta Q^2\mu^{2Q} - \\ &\big(2w(Q-1) + \beta(Q-1)\big)\alpha Q\mu^Q - w(Q-1)(Q-2)\Big]. \end{aligned} \tag{17}$$

Again, condition 8 is satisfied if the term in the square brackets is negative. Notice that this term is a quadratic polynomial in $s = \alpha Q \mu^Q$,

$$-\beta s^2 - \big(2w(Q-1) + \beta(Q-1)\big)s - w(Q-1)(Q-2). \tag{18}$$

. This polynomial has two positive real roots: the discriminant $\Delta = 4w^2(Q-1)^2 + \beta^2(Q-1) + 4w\beta(Q-1) > 0$. Moreover it is negative and increasing at $s = 0$ and take a maximum at a positive value.

Therefore, we have the following proposition

**Proposition 1** *Let $w(\mu)$ be the fitness function defined by Equation 1 and the genotypic* *fitness $W(\mu, \sigma)$ defined by Equation 2. Then*

$$\frac{\partial^2 \ln W(\mu,\sigma)}{\partial\mu\partial\sigma} > 0$$

*if and only if one of the following properties is satisfied.*

- *$Q = 2$, $\beta > 0$, and $\mu^2 > (2w + \beta)/(2\alpha\beta)$,*
- *$Q > 2$, $\beta = 0$, and $\mu^Q < (Q - 2)/(2\alpha Q)$,*
- *$Q > 2$, $\beta > 0$, and $\mu \in I_1 \cup I_2$, where the intervals $I_1 = ]0, \mu_1[$ and $I_2 = ]\mu_2, +\infty[$, and $\mu_1, \mu_2$ are the two positive roots of the polynomial (Eq. 18).*

As shown on Figure 1A, these inequalities are satisfied **when the logarithmic fitness plateaus** (else the optimum $\ln(1)$, else the minimum $\ln(\beta)$, as the fitness curvature tends to zero, Fig. 1C). The exact values of $\mu$ for which phenotypic noise promotes adaptive evolution depends on the difference between the relative fitness gradient $\frac{\partial \ln w(\mu)}{\partial\mu}$ and the relative change of the curvature $\frac{\partial \ln w''(\mu)}{\partial\mu}$, dictated by the difference $w(\mu)w'''(\mu) - w'(\mu)w''(\mu)$ (Fig. 1B).

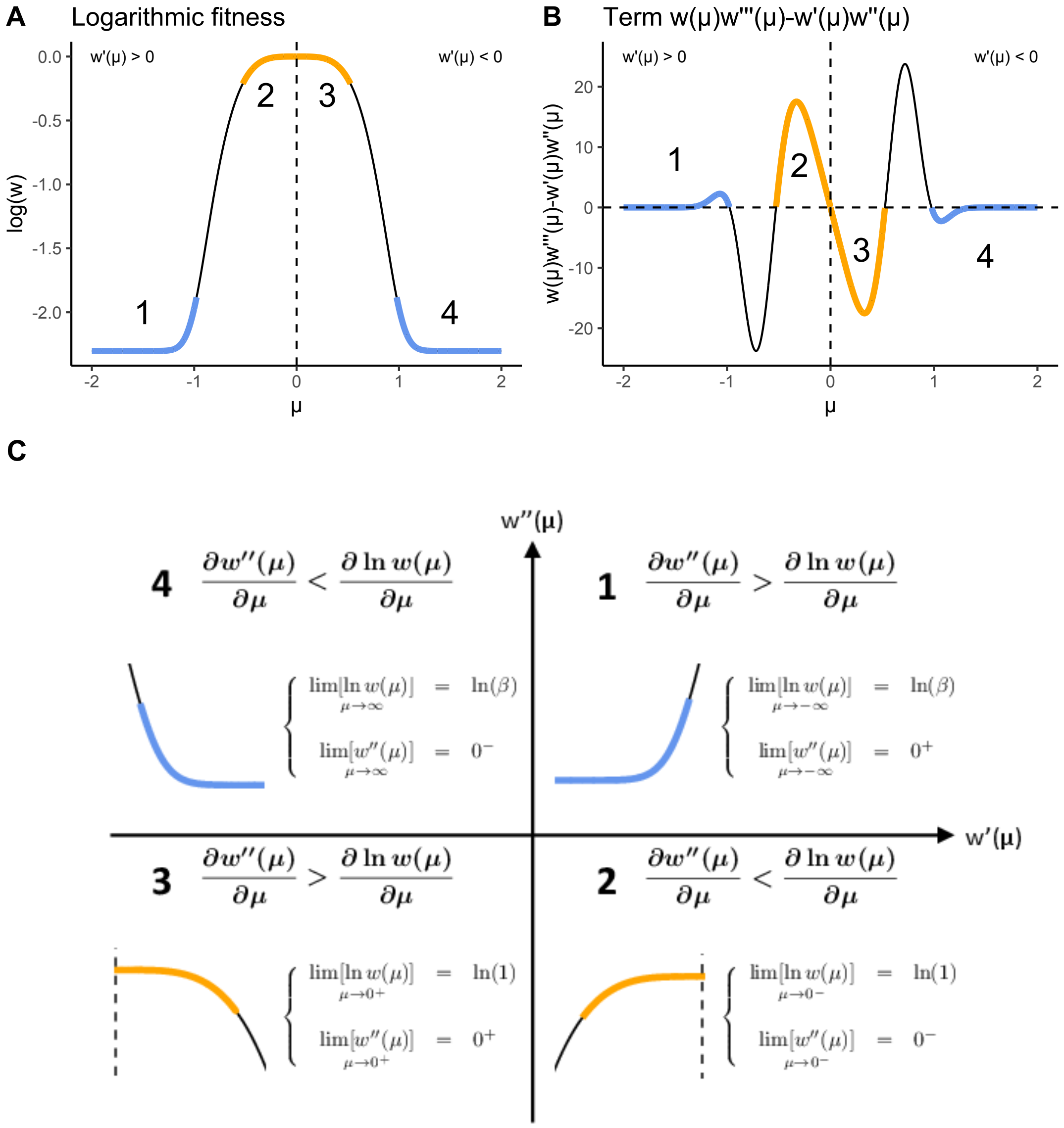


Figure 1 – **Phenotypic noise increases the gradient of relative genotypic fitness when the logarithmic fitness plateaus. A.** Logarithmic fitness $\ln w(z)$ where $\alpha = 3.125$, $\beta = 0.1$ and $Q = 4$. When $\mu$ is in the blue and orange regions (numbered from 1 to 4), phenotypic noise increases the gradient of relative genotypic fitness. **B.** Term $w(\mu)w'''(\mu) - w'(\mu)w''(\mu)$ depending on $\mu$ (Eq. 8). When $\mu$ is in the blue and orange regions (numbered from 1 to 4), phenotypic noise increases the gradient of relative genotypic fitness. **C.** Four cases corresponding to regions 1, 2, 3 and 4. In all cases, the analysis of the term $w(\mu)w'''(\mu) - w'(\mu)w''(\mu)$ shows that phenotypic noise increases the gradient of relative genotypic fitness when the logarithmic fitness tends to a constant $>$ 0 and when the fitness curvature tends to 0, *i.e.* when the logarithmic fitness plateaus.

### 3 For multiple characters under selection, the best phenotypic noise configuration is fully correlated and aligned with the fitness optimum

Let us consider the organism with genotype $\{\boldsymbol{\mu}, \boldsymbol{\sigma}, \boldsymbol{\theta}\}$ (see main manuscript) in a $n$-dimensional phenotypic space, sitting at a certain distance of the fitness optimum $\boldsymbol{z_{opt}} = \mathbf{0}$ of the fitness function $w(\boldsymbol{z}) = (1-\beta)e^{-\alpha\|\boldsymbol{z}\|^Q} + \beta$ and under stabilizing selection (with $\|\boldsymbol{\mu}\| > |z_{\text{infl}}|$, see main manuscript). We describe the phenotypic noise of this organism by a multivariate normal distribution $\mathcal{N}_n(\boldsymbol{\mu}, \boldsymbol{\Sigma})$, $\boldsymbol{\Sigma}$ being the $n \times n$ covariance matrix built from $\boldsymbol{\sigma}$ and $\boldsymbol{\theta}$ (see main manuscript). This multivariate normal distribution can be represented by an hyper-ellipse in $\mathbb{R}^n$, as shown in Figure 2 for two phenotypic characters ($n = 2$).

We now define an orthonormal set of vectors $\boldsymbol{v_1}, ..., \boldsymbol{v_n}$, forming a new basis with origin at $\boldsymbol{\mu}$. We align the direction of the first element $\boldsymbol{v_1}$ towards the fitness optimum $\boldsymbol{z_{opt}}$ (Fig. 2A). Along the axis $\mu + a\boldsymbol{v_1}$, $a$ being a scalar, the organism $\{\boldsymbol{\mu}, \boldsymbol{\sigma}, \boldsymbol{\theta}\}$ experiences a convex fitness if $\|\boldsymbol{\mu}\| > |z_{\text{infl}}|$. Along all other axes $\mu + a\boldsymbol{v_i}$, the organism experiences a concave fitness, and $\mu$ is a local fitness optimum for all values of $a$ (Fig. 2B). We denote by $\boldsymbol{V}$ the matrix formed by the vector column $\boldsymbol{v_i}$,

$$\boldsymbol{V} = \left(\boldsymbol{v_1}\middle|\boldsymbol{v_2}\middle|...\right) \tag{19}$$

The goal here is to find the phenotypic noise configuration $\boldsymbol{\Sigma}$ that maximizes the genotypic fitness

$$W(\boldsymbol{\mu}, \boldsymbol{\Sigma}) = \int_{\mathbb{R}^n} p(\boldsymbol{z}|\boldsymbol{\mu}, \boldsymbol{\Sigma}) w(\boldsymbol{z}) d\boldsymbol{z}. \tag{20}$$

We first make a change of variables to center the phenotype around $\boldsymbol{\mu}$: $\boldsymbol{z} = \boldsymbol{\mu} + \boldsymbol{\epsilon}$, to obtain

$$W(\boldsymbol{\mu}, \boldsymbol{\Sigma}) = \int_{\mathbb{R}^n} p(\boldsymbol{\epsilon}|\mathbf{0}, \boldsymbol{\Sigma}) w(\boldsymbol{\mu} + \boldsymbol{\epsilon}) d\boldsymbol{\epsilon}. \tag{21}$$

If $\|\boldsymbol{\mu}\| > |z_{\text{infl}}|$, phenotypic noise correlated with $\boldsymbol{v_1}$ increases the genotypic fitness, and phenotypic noise correlated with any other directions $\boldsymbol{v_i}, i > 1$ decreases the genotypic fitness (Fig. 2B): For any covariance matrix $\boldsymbol{\Sigma}$ and for any $\boldsymbol{\epsilon} \sim \mathcal{N}_n(\boldsymbol{0}, \boldsymbol{\Sigma})$, the fitness $w(\boldsymbol{\mu}+\boldsymbol{\epsilon})$ of the phenotype $\boldsymbol{z} = \boldsymbol{\mu}+\boldsymbol{\epsilon}$ is always lower or equal to the fitness of its projection along the axis defined by $\boldsymbol{v_1}$ (*i.e.*, the distance to the fitness optimum is shorter after the projection). Thus:

$$\int_{\mathbb{R}^n} p(\boldsymbol{\epsilon}|\boldsymbol{0}, \boldsymbol{\Sigma}) w(\boldsymbol{\mu} + \boldsymbol{\epsilon}) d\boldsymbol{\epsilon} \leq \int_{\mathbb{R}^n} p(\boldsymbol{\epsilon}|\boldsymbol{0}, \boldsymbol{\Sigma}) w(\boldsymbol{\mu} + \boldsymbol{v_1}^T \boldsymbol{\epsilon} \boldsymbol{v_1}) d\boldsymbol{\epsilon}. \tag{22}$$

We then express $\boldsymbol{\epsilon}$ in the basis $\boldsymbol{V}$ by making the following variable change: $\boldsymbol{s} = \boldsymbol{V}^T\boldsymbol{\epsilon}$. Consequently, $\boldsymbol{\epsilon} = \boldsymbol{V}\boldsymbol{s}$, such that:

$$\begin{aligned} \boldsymbol{v_1}^T \boldsymbol{\epsilon} \boldsymbol{v_1} &= \boldsymbol{v_1}^T \boldsymbol{V} \boldsymbol{s} \boldsymbol{v_1} \\ &= s_1 \boldsymbol{v_1}. \end{aligned} \tag{23}$$

($s_1$ is the first coefficient of the vector $\boldsymbol{s}$). We can rewrite the right term of the Equation 22 as follows:

$$\int_{\mathbb{R}^n} p(\boldsymbol{V}\boldsymbol{s}|\boldsymbol{0}, \boldsymbol{\Sigma}) w(\boldsymbol{\mu} + s_1 \boldsymbol{v_1}) ds_1, ..., ds_n. \tag{24}$$

The term $w(\boldsymbol{\mu} + s_1\boldsymbol{v_1})$ depends only on $s_1$, so we can re-organize the integral,

$$\int_{\mathbb{R}} w(\boldsymbol{\mu} + s_1 \boldsymbol{v_1}) \left[ \int_{\mathbb{R}^{n-1}} p(\boldsymbol{V}\boldsymbol{s}|\boldsymbol{0}, \boldsymbol{\Sigma}) ds_2, ..., ds_n \right] ds_1. \tag{25}$$

The probability density function $p(\boldsymbol{V}\boldsymbol{s}|\boldsymbol{0}, \boldsymbol{\Sigma})$ from Equation 25 is the same as $p(\boldsymbol{s}|\boldsymbol{0}, \boldsymbol{V}^T\boldsymbol{\Sigma}\boldsymbol{V})$ (by a change of variable). The inside integral in Equation 25,

$$\int_{\mathbb{R}^{n-1}} p(\boldsymbol{s}|\boldsymbol{0}, \boldsymbol{V}^T \boldsymbol{\Sigma} \boldsymbol{V}) ds_2, ..., ds_n, \tag{26}$$

describes the marginal density of $s_1$, and follows the univariate normal law:

$$s_1 \sim \mathcal{N}\Big(0, \left[\boldsymbol{V}^T \boldsymbol{\Sigma} \boldsymbol{V}\right]_{1,1}\Big) \tag{27}$$

the subscript "1, 1" denoting the coefficient of the first raw and first column.

Therefore, the integral 25 simplifies to

$$\int_{\mathbb{R}} w(\boldsymbol{\mu} + s_1 \boldsymbol{v_1}) p(s_1 | 0, \left[\boldsymbol{V}^T \boldsymbol{\Sigma} \boldsymbol{V}\right]_{1,1}) ds_1. \tag{28}$$

Equation 28 is the one dimensional genotypic fitness $W(\boldsymbol{\mu}, \left[\boldsymbol{V}^T \boldsymbol{\Sigma} \boldsymbol{V}\right]_{1,1})$, taken along the axis spanned by $v_1$. Along this axis, there is an optimal phenotypic noise amplitude $\left[\boldsymbol{V}^T \boldsymbol{\Sigma} \boldsymbol{V}\right]_{1,1} = \sigma^2_{opt}$ that maximizes the genotypic fitness. The $n$-dimensional genotypic fitness can be maximized by the 1-dimensional genotypic fitness

$$\begin{aligned} W(\boldsymbol{\mu}, \boldsymbol{\Sigma}) &\leq \int_{\mathbb{R}} w(z) p(z | \|\boldsymbol{\mu}\|, \left[\boldsymbol{V}^T \boldsymbol{\Sigma} \boldsymbol{V}\right]_{1,1}) dz \\ &= W(\|\boldsymbol{\mu}\|, \left[\boldsymbol{V}^T \boldsymbol{\Sigma} \boldsymbol{V}\right]_{1,1}) \\ &\leq W(\|\boldsymbol{\mu}\|, \sigma^2_{opt}). \end{aligned}$$

Therefore, the $n$-dimensional genotypic fitness reaches its maximum when all the phenotypic noise is concentrated along the axis aligned with the direction to the fitness optimum. By expressing the covariance matrix $\boldsymbol{\Sigma}$ as an eigenvalue factorization, $\boldsymbol{\Sigma} = \boldsymbol{U} \boldsymbol{D} \boldsymbol{U}^T$, with $\boldsymbol{D} = \mathrm{diag}(\boldsymbol{\sigma}^2)$, we obtain:

$$\boldsymbol{V}^T \boldsymbol{\Sigma} \boldsymbol{V} = \boldsymbol{V}^T \boldsymbol{U} \boldsymbol{D} \boldsymbol{U}^T \boldsymbol{V}. \tag{29}$$

The maximum genotypic fitness is reached when the first eigenvector $\boldsymbol{u_1}$ is aligned with $\boldsymbol{v_1}$ and $\boldsymbol{\sigma}^2 = \{\sigma^2_{opt}, 0, ..., 0\}$ (Fig. 2A).

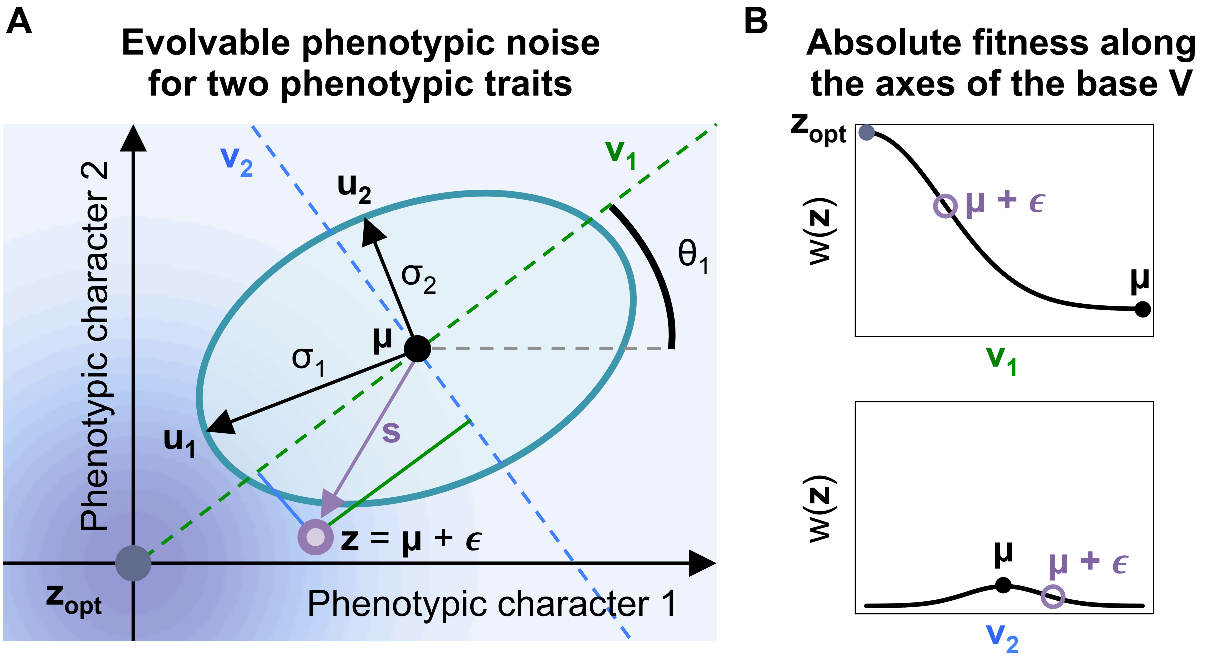


Figure 2 – **Anisotropic and correlated phenotypic noise for two phenotypic characters under selection.** **A.** The phenotypic noise distribution of an organism with genotype $\{\boldsymbol{\mu}, \boldsymbol{\sigma}, \boldsymbol{\theta}\}$ is defined by a multivariate normal distribution with mean $\boldsymbol{\mu}$ (black dot), phenotypic noise amplitudes $\sigma_1$ and $\sigma_2$ (black arrows) along axes $\boldsymbol{u_1}$ and $\boldsymbol{u_2}$, and a parameter of correlation $\theta_1$ (black angle), defining a rotation of the basis $\boldsymbol{U} = (\boldsymbol{u_1}, \boldsymbol{u_2})$. A phenotype $\boldsymbol{z}$ (purple dot) is generated from the multivariate normal distribution by drawing a random vector $\boldsymbol{\epsilon} \sim \mathcal{N}_n(\mathbf{0}, \boldsymbol{\Sigma})$ (with $\boldsymbol{\Sigma}$ the covariance matrix built from $\boldsymbol{\sigma}$ and $\boldsymbol{\theta}$), such that $\boldsymbol{z} = \boldsymbol{\mu} + \boldsymbol{\epsilon}$. The contribution of $\boldsymbol{\epsilon}$ on each axis $\boldsymbol{v_1}$ (green solid segment) and $\boldsymbol{v_2}$ (blue solid segment) of the basis $\boldsymbol{V}$, where $\boldsymbol{v_1}$ is aligned with the fitness optimum $\boldsymbol{z_{opt}}$, is represented by the vector $\boldsymbol{s} = (s_1, s_2)^T$ (purple arrow). The fitness landscape is represented by a gradient of blue centered on the fitness optimum $\boldsymbol{z_{opt}}$ (deep blue dot). **B.** Fitness along axes of the basis $\boldsymbol{V} = (\boldsymbol{v_1}, \boldsymbol{v_2})$. Along axis $\boldsymbol{v_1}$, directed towards the fitness optimum $\boldsymbol{z_{opt}}$, the organism experiences a convex absolute fitness (if $\|\boldsymbol{\mu}\| > |z_{\text{infl}}|$). Along axis $\boldsymbol{v_2}$, orthogonal to $\boldsymbol{v_1}$, the organism experiences a concave absolute fitness, sitting on a local fitness optimum.

### References

Lande, R. (1979). Quantitative genetic analysis of multivariate evolution, applied to brain: body size allometry. *Evolution*, 33(1Part2):402–416.

Martin, G. (2014). Fisher's geometrical model emerges as a property of complex integrated phenotypic networks. *Genetics*, 197(1):237–255.

Paenke, I., Sendhoff, B., and Kawecki, T. J. (2007). Influence of plasticity and learning on evolution under directional selection. *The American Naturalist*, 170(2):E47–E58.

Slatkin, M. and Lande, R. (1976). Niche width in a fluctuating environment-density independent model. *The American Naturalist*, 110(971):31–55.
